# Contributions of human-caused climate change and individual emitters to global coral bleaching

**DOI:** 10.64898/2026.09.08.750055

**Authors:** Maximilian Kotz, Puja Pande, Rory Gibb, Andreas Schwarz Meyer, Olivia Bates, Christopher Callahan, Markus Donat, Christopher Trisos

**Affiliations:** Barcelona Supercomputing Centre, Barcelona; Potsdam Institute for Climate Impact Research, Potsdam; Centre for Biodiversity & Conservation Science, University of Queensland, Brisbane; Climate Risk Lab, African Climate and Development Initiative, University of Cape Town; Department of Genetics, Evolution and Environment, University College London; O’Neill School of Public & Environmental Affairs, Indiana University, Bloomington; Institució Catalana de Recerca i Estudis Avançats (ICREA), Barcelona; African Synthesis Centre for Climate Change Environment and Development (ASCEND), African Climate and Development Initiative, University of Cape Town

**Author notes:** These authors share first author status author (Maximilian Kotz, Puja Pande).

## Abstract

Coral bleaching is among the most infamous climate impacts^1,2^. Yet the extent of bleaching due to human-caused climate change, as opposed to natural El Niño variability, and the contributions of individual countries and carbon majors (fossil fuel and cement producers) remains unquantified. Here, we leverage coral monitoring globally across 4,479 sites from 1985–2024^3,4^ to link specific emitters to marine heat stress and bleaching. For events with >10% of corals bleached at a site, we find that 99% (95% CI: 88–100%) were attributable to human-caused climate change. Even during major El Niños, 97–100% of bleaching events were attributable to human-caused climate change and the extent of damage increases over time from 23% of sites bleached (95% CI: 3–35%) during 1998 to 71% of sites (95% CI: 37–85%) during 2023–2024. Emissions of the United States, the largest historical emitter, contributed to 51% (95% CI: 39–65%) of bleaching events. Emissions linked to Chevron, Saudi Aramco, Gazprom, and ExxonMobil each contributed to 14% (95% CI: 11–19%) of bleaching events—roughly double the contribution of the 39 countries of the Alliance for Small Island States. It is virtually certain emissions of China, the United States, and Indonesia caused bleaching in their coastal waters. Strong inequalities exist where countries with high socioeconomic dependence on coral reefs experienced 54% of bleaching yet contributed only 5% to bleaching. As global warming rapidly nears levels catastrophic for warm-water corals^1,5^, our results show the massive contribution from anthropogenic emissions to historical coral beaching and help fill an important evidentiary gap on accountability for non-economic loss and damage from climate change.

## Introduction

Warm-water coral reefs house approximately one-quarter of marine biodiversity^6^, support the livelihoods of hundreds of millions of people^7–9^, and have suffered significant adverse impacts from marine heat extremes—especially during recent El Niño events (e.g., 2023–2024)^2,10–12^. Heat stress causes corals to expel the symbiotic *zooxanthellae* algae on which they depend^13^, leading to loss of colour (hereafter, bleaching) and in many cases post-bleaching mortality^14^. Concern about potentially catastrophic impacts for coral reefs from global warming became widespread after the 1998 El Niño event when high sea surface temperatures (SSTs) led to the first global coral bleaching episode^15^. It is now unequivocal that anthropogenic activities, principally through greenhouse gas emissions from fossil fuel combustion, are warming the ocean^16^, including in coral regions^17,18^. Yet we still lack quantification of how much coral bleaching globally is due to human-caused climate change (as opposed to natural climate variability such as El Niño^19^), whether human-caused climate change’s contribution has changed over time, and how much bleaching is attributable to the emissions of specific actors, such as sovereign states or companies.

Improved understanding of the global human impact on coral reefs requires determining how much bleaching has been driven by anthropogenic climate change. Extending this attribution to individual states and companies specifically is important for understanding inequalities among contributions to, and harms from, bleaching impacts. The quantification of impacts on coral reefs from human-caused climate change generally, and from individual emitters specifically, is timely to advance loss and damage policy discussions, given a predominant focus to date on economic over non-economic loss and damage^20,21^. It is also significant for informing climate justice processes more generally, including legal liability, given many tropical countries and communities directly dependent on coral reefs (e.g., Small Island Developing States) have contributed less to the emissions driving global warming^22^.

The lack of an end-to-end quantification of the causal chain from specific emitters to observed coral bleaching partly reflects important methodological challenges for attribution of coral reef bleaching (as well as for biodiversity impact attribution more generally)^23^. For example, coral reef communities, and their responses to heat, are heterogeneous at much finer spatial resolutions^24^ than most global climate model simulations, rendering studies attributing marine heat-waves^25^ and rising SSTs within large-scale ocean regions^18^ ill-equipped for attribution of bleaching impacts. Furthermore, the El Niño Southern Oscillation is the dominant determinant of the timing of recent coral heat exposure^26^, but attribution methodologies using climate models do not reproduce the observed timing of this natural climate variability (e.g. from the Detection Attribution Model Intercomparison Project^27^).

Here, we combine advances across causal inference frameworks, global coral reef monitoring data, and climate change counterfactual methods to quantify the contribution to coral heat exposure and bleaching from anthropogenic emissions in general, as well as for specific states and major oil, gas, coal, and cement producers (hereafter, ‘carbon majors’) (see Methods). Using a synthesis of global coral bleaching data^3^ supplemented with more recent observations^28^ we obtain 4,479 coral sites with population-level bleaching data covering all El Niño events from 1998–2024 across all major warm-water coral regions (Figure 1a). We calculate historical heat exposure for these sites using satellite-derived SST observations at 5km resolution extending back to 1985^29^. Following decades of coral reef science, we use the well-established degree heating weeks (DHWs) metric to measure the intensity of heat exposure for corals^10,12,29^, defined as the cumulative 12-week sum of degrees of SST that are 1°C above the maximum monthly mean SST climatology at a given reef site. Next, using the subset of 1,626 reef sites with repeat bleaching observations, we leverage climate econometrics^30,31^ and ecology^23^ methods for inferring causal relationships between climate hazards and impacts to develop a panel regression model that isolates the effect of observed DHWs on coral bleaching from other confounding factors. Nonparametric controls in the model (that is, fixed effects) account for regional differences in seasonality, for time-invariant site-specific unobserved confounders (e.g., depth or distance from shore), and for confounders common across sites in a given year (e.g., influence of El Niño events). We then apply this model to estimate coral bleaching for historical time series of DHWs and for counterfactual time series with either all anthropogenic emissions or only specific emission sources removed, accounting for both statistical and climatological uncertainty. We generate counterfactuals of DHW that retain the high spatial resolution of observed SST data and preserve the observed timing of El Niño events associated with coral bleaching, while removing the warming driven by anthropogenic emissions. To achieve this, we use the Finite amplitude Impulse Response (FaIR) simple climate model^32^ to generate counterfactuals of global mean surface temperature (GMST) excluding specific emissions (e.g., from a carbon major)^33^, and then we use pattern-scaling relationships derived directly from observed daily SSTs and observed GMST^34,35^ to map counterfactual GMST to counterfactual DHW values for coral reef sites. We account for uncertainty in transient climate sensitivity by sampling the full set of 841 posterior ensemble members available for FaIR model parametrisation^32^, and we also quantify climate uncertainty in the pattern scaling of SSTs using a bootstrap approach designed to accurately reflect uncertainty due to natural climate variability when using temperature observations directly for attribution^34^. This avoids the need to use SST data from global climate models which would require bias-adjustment to reflect SSTs at coral reef sites^36^, would not preserve the timing of El Niño events, and would also risk inclusion of significant biases in global climate model SST trends in important ocean regions for coral reefs, such as the tropical Indo-Pacific^37^.

**Fig. 1.**
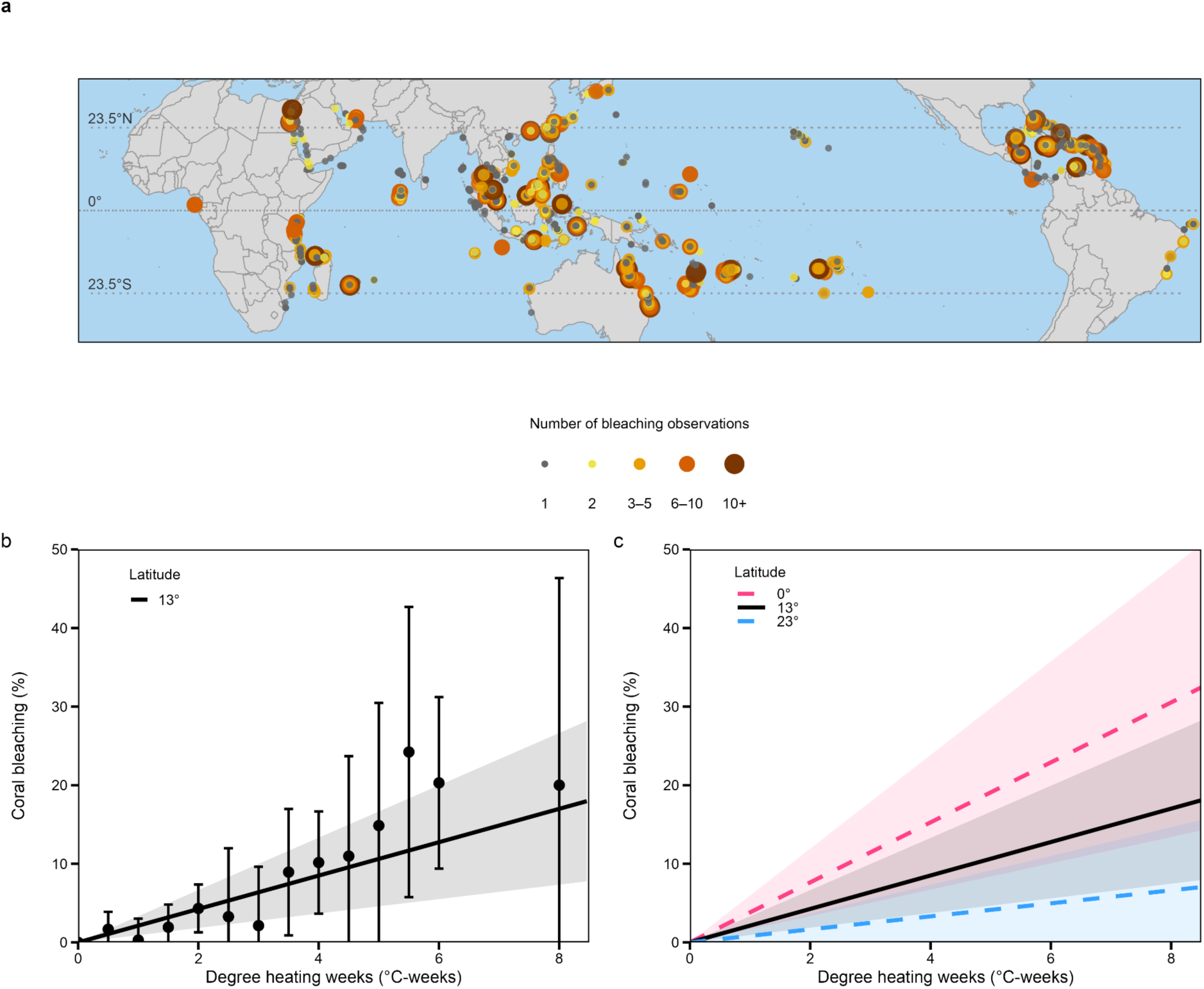
Coral bleaching observations and empirical estimates of bleaching–marine heat relationships. **a**, The location and number of observations from coral reef bleaching surveys from 1998–2024 used in this study. **b**, The estimated relationship between Degree Heating Weeks and percentage coral bleaching, based on a linear fixed-effects model (shading indicates 95% confidence intervals accounting for spatial autocorrelation up to 200km). Estimates from a non-parametric binned fixed-effects model (point estimates and 95% confidence intervals) are also shown, revealing limited non-linearities and high consistency with the linear model. Estimates are shown for the median absolute latitude observed in the sample (13°N or S). **c**, The effect of DHW on bleaching varies by absolute latitude, with stronger effects at lower latitudes. Estimates of percent bleaching for a given level of degree heating weeks are shown at the equator (0°), the median absolute latitude observed in the sample (13°N or S) and at the edge of the tropics (23°N or S). Shading indicates 95% confidence intervals.

### A robust signal of heat on coral bleaching

We estimate the effect of DHW on bleaching as a linear response in a panel regression causal-inference model, finding a significant effect of increasing DHWs on bleaching (Fig. 1b): For sites located at 13° latitude (the mean latitude of monitored sites) an additional 4 °C-week exposure leads to an increase in the percentage of coral colonies bleaching at a site of 8.5% (95% CI: 3.7–13.3%). We test whether this effect of DHW on bleaching varies based on potential moderators and find the relationship does not vary significantly with a site’s Marine Protected Area status, an indicator of local human pressures other than climate change, consistent with findings that protected areas do not buffer corals from heat-driven bleaching^2,38^ (Supplementary Table 1). We also find the bleaching–DHW relationship does not vary significantly with depth (0–20m depth range for sites in this study), consistent with prior global studies^39^; nor with turbidity, noting prior work suggests moderately turbid conditions may reduce bleaching although this effect may be weak and present only for low levels of heat stress^40,41^; nor with historical rates of sea surface warming (a test of potential local adaptation to heat stress in faster warming environments^42^) (Supplementary Table 1). However, the effect of DHW on bleaching is significantly higher closer to the equator and at sites that have historically experienced lower SST variability (Fig. 1c, Supplementary Table 1). These effects likely reflect a combination of historical adaptation to higher SST variability as well as broad biogeographic differences in the diversity and composition of coral species^39,43^. We prefer a model including only the moderating effect of absolute latitude on the bleaching–DHW relationship because of the high collinearity between latitude and SST variability (*r =* 0.7), and the very similar patterns of coral bleaching sensitivity to DHW when using either SST variability or latitude as a moderator (Extended Data Fig. 1). For the same level of 4°C-weeks additional heat exposure, coral reef sites at the equator are estimated to experience nearly twice as much bleaching (15.3%, 95% CI: 6.8–23.9%) as sites at the mean latitude of monitored sites (13° N or S) (8.5%, 95% CI: 3.7–13.3%; Figure 1c).

Additional sensitivity analyses reinforce that this bleaching–DHW relationship is robust. Specifically, a fully non-parametric model of DHW impacts on bleaching with binned DHW exposure values that flexibly allow for non-linearities in the bleaching response to DHW is largely consistent with the linear model (Fig. 1b); higher-order polynomial effects of DHW do not improve on the linear model (Supplementary Table 2); and the response of bleaching to DHW is similar when using alternative methods explicitly modeling the zero-inflated nature of the bleaching data (Supplementary Figs. 1 and 2, see Methods). Results are also robust when adjusting DHW to be accumulated over fewer weeks and a lower exceedance of maximum monthly mean SST climatology, as suggested for some bleaching forecasts^44^ (Supplementary Table 3). Assessments of uncertainty in all models account for spatial autocorrelation of model residuals which decay to approximately zero beyond distances of 200km with results robust to alternative error specifications (Supplementary Fig. 3 and Supplementary Table 4).

**Fig. 2.**
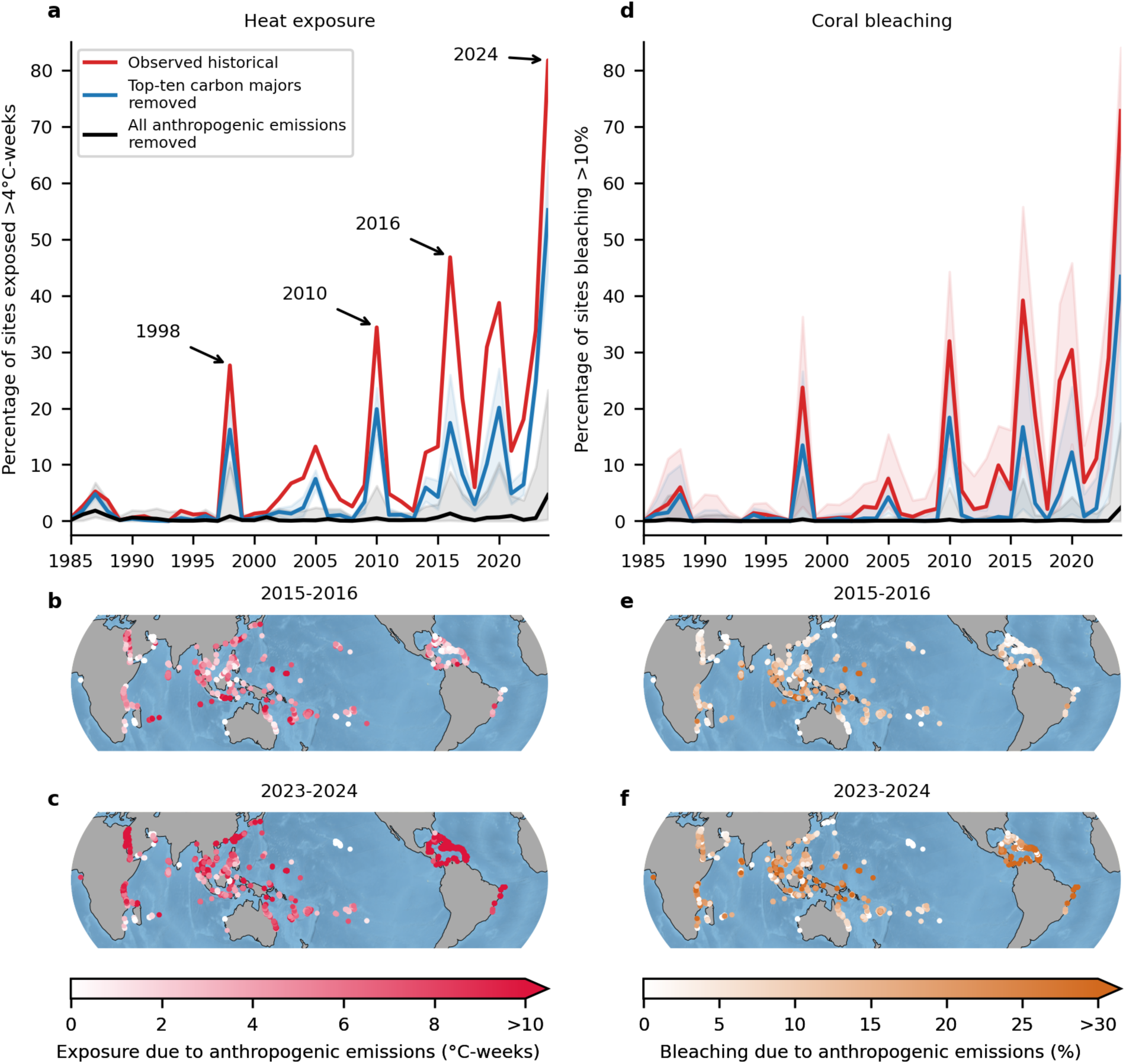
Historical heat exposure and bleaching of coral reefs attributable to human-caused climate change. **a,** Percentage of coral sites in the global sample exposed to greater than four degree heating weeks under observed sea-surface temperatures from 1985–2024 (red); counterfactual climates with all anthropogenic emissions removed (black); and counterfactual climates with the emissions of the top ten emitting carbon majors removed (blue). Solid lines show the median and shaded areas the 95% (extremely likely) confidence intervals. Years of major global bleaching are highlighted. **b, c,** Additional heat exposure during the 2016 and 2024 El Niño events that was attributable to anthropogenic emissions. **d,** Percentage of coral reef sites in the global sample estimated to have more than 10% of coral colonies bleached for observed sea-surface temperatures (red); counterfactual climates with all anthropogenic emissions removed (black); and the emissions of the top ten emitting carbon majors removed (blue). Solid lines show the median and shaded areas the 95% confidence intervals. **e, f,** Additional coral bleaching during the 2016 and 2024 El Niño events that was attributable to anthropogenic emissions.

### Massive extent of bleaching attributable to human-caused climate change

Heat exposure for corals has become more frequent and more intense since the 1980s with 82% of coral reef sites during the 2024 El Niño experiencing greater than 4°C-weeks of heat exposure—a threshold used to issue bleaching alerts^29^ (Fig. 2a).

Comparing the observed historical climate to counterfactual climates with anthropogenic climate forcing removed, we find that coral reef heat exposure and bleaching would have been minimal were it not for human-caused climate change (Fig. 2, Extended Data Figs. 2 and 3). Over the period 1985–2024, human-caused climate change was responsible for 97% of coral reef heat exposure events above 4 °C-weeks (95% [extremely likely] CI range: 87–99%), and for 99% of bleaching events (95% CI: 88–100%) where over 10% of corals at a site bleached (Fig. 2a,d, Supplementary Table 5). These percentages represent 20,042 heat exposure events (95% CI: 17,880–20,465) and 15,345 bleaching events (95% CI: 4,556–25,345) attributable to human-caused climate change from 1985–2024 across the 4,475 sites in our global sample. Ten percent bleaching is widely used as a threshold for describing the transition from light to moderate levels of bleaching within coral reef research^2,11^. The confidence in the contribution of human-caused climate change to bleaching rises with higher bleaching thresholds such that for bleaching events where more than 20% of corals bleached, 99% of these events were due to human-caused climate change and with a narrowed confidence interval (95 % CI: 94–100%) (Supplementary Table 6, Extended Data Fig. 3). This extent of damage attributable to human-caused climate change is relatively unprecedented for a pan-tropical or global impact on humans or ecosystems, but aligns with recent risk assessments of warm-water coral reefs being Earth’s first climate ‘tipping point’^45^.

**Fig. 3.**
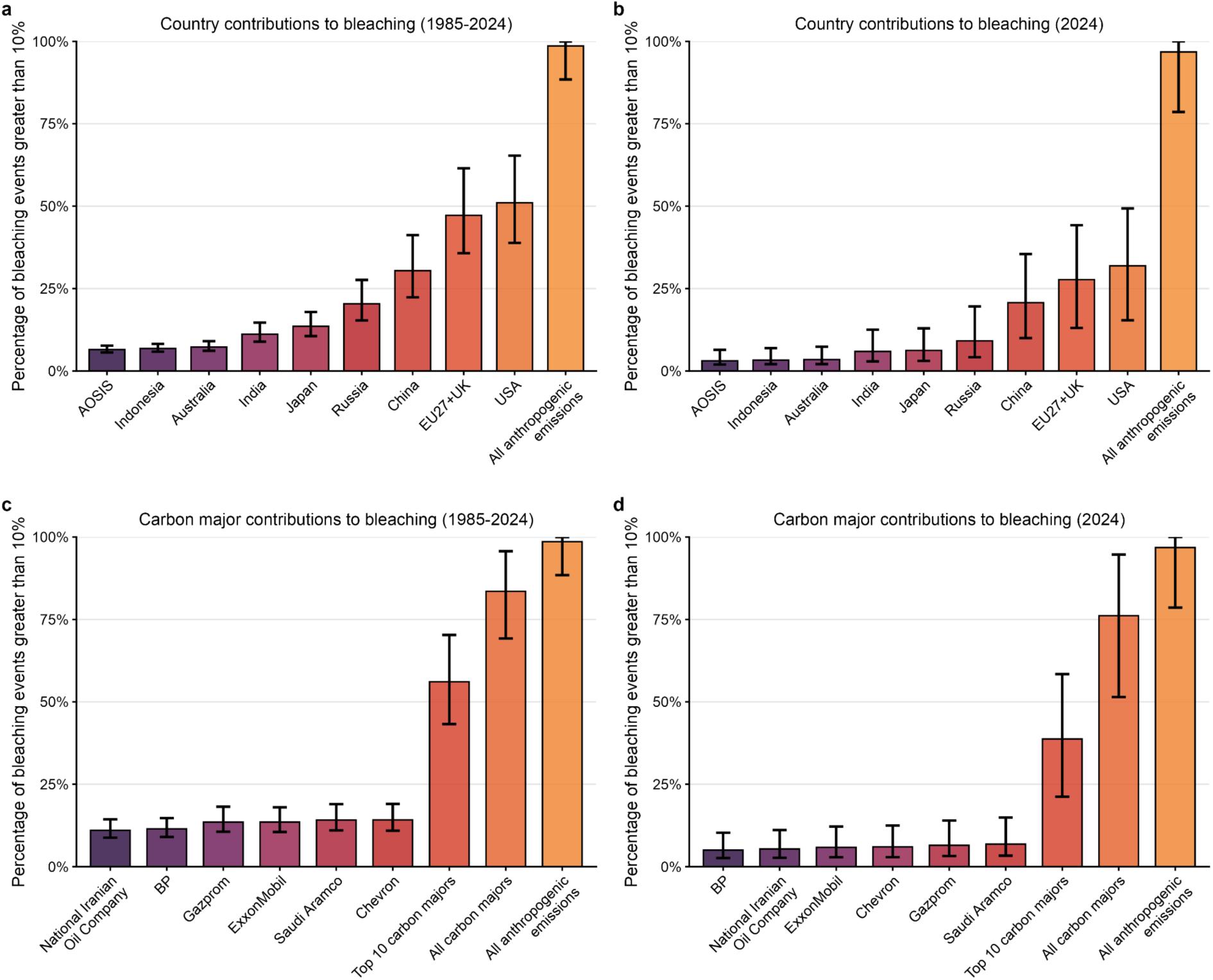
Contributions to coral bleaching from marine heat by states and carbon majors. **a,** The percentage of coral reef bleaching events of over 10% from 1985–2024 which would not have occurred without the territorial emissions of individual states or regional blocs. **b,** The percentage of coral reef bleaching events of over 10% during the 2023–2024 El Niño which would not have occurred without the territorial emissions of individual states or regional blocs. **c,d**, as for panels a and b, but showing contributions from the emissions of major emitting companies (‘carbon majors’). Bar plots are coloured based on the size of the contribution to coral bleaching and error bars show 95% confidence intervals. “All carbon majors” refers to the emissions of 172 investor or state-owned fossil fuel companies in the Carbon Majors database over the period 1854–2024. AOSIS is the 39 countries participating in the Alliance of Small Island States and EU27+UK refers to the 27 countries of the European Union and the United Kingdom. See Supplementary Tables 5–8 for values for individual states and companies.

Taking a finer temporal lens enables further unpacking of the causes of heat-driven coral bleaching during El Niño years when SSTs are elevated in large parts of the tropical oceans (Fig. 2a,d). Using climate counterfactuals that preserve the timing of observed natural climate variability in SSTs, shows human-caused climate change has been the overwhelming contributor to coral bleaching even during El Niño events: without human-caused climate change the percentage of coral reef sites experiencing greater than 4 °C-weeks of heat stress and more than 10% coral bleaching would have remained near zero during all El Niño years since 1985 (Fig. 2a,d, Extended Data Figs. 4 and 5). Taking the difference between the observed and counterfactual climates allows us to quantify the specific anthropogenic contribution to levels of coral reef heat exposure and bleaching within individual El Niño years (Fig. 2b,c,e,f). For example, due to human-caused climate change, we estimate that during the 2023–2024 El Niño 80% of coral reef sites (95% CI: 60–85%) were exposed to greater than 4 °C-weeks of heat stress and 49% of sites (95% CI: 44–50%) were exposed to greater than 8 °C-weeks of heat stress—a threshold for risk of coral mortality^29^—with this heat exposure especially strong in the Caribbean (Fig. 2c; Extended Data Fig. 4). We estimate that 97% of events (95% CI: 79–100%) with over 10% coral bleaching during the 2023–2024 El Niño were attributable to human-caused climate change, affecting 71% of reef sites (95% CI: 37–85%), and with events of over 20% coral bleaching affecting 39% of sites (95% CI: 9–65%) (Fig. 2f; Extended Data Fig. 5).

The contribution of human-caused climate change has increased the extent of global coral reef heat exposure and bleaching with every major El Niño since 1998 (Extended Data Figs. 6 and 7). Human-caused climate change was responsible for 99% of events with more than 10% coral bleaching, affecting 23% of reef sites, during the 1998 El Niño; 99% of events affecting 31% of sites during 2010; 100% of events affecting 39% of sites during 2015–2016; and 97% of events affecting 71% of sites during the 2023–2024 El Niño. This contribution of human-caused climate change to increasing coral bleaching with successive El Niño events is robust across multiple thresholds used to define heat exposure and bleaching (Extended Data Figs. 6 and 7). These results demonstrate the unequivocal role of human-caused climate change in driving virtually all heat exposure and bleaching of coral reefs from 1985–2024, vastly outweighing the role of natural climate variability from El Niño.

### Coral bleaching from countries and carbon majors

Counterfactual scenarios that exclude emissions from specific sources provide a means to quantify contributions to bleaching. These scenarios follow the logic of ‘but for’ causation^33,46^ to ask: would the prevalence of coral bleaching above a given threshold have been lower if the emissions of a specific country or carbon major had not occurred? For example, Figure 2a and 2d show heat exposure and bleaching under a counterfactual scenario without the emissions of the top ten carbon majors (red vs. blue).

Without the emissions of the United States, the country with the largest cumulative territorial emissions since 1850, 51% of events with over 10% coral bleaching at a site would not have occurred over the period 1985–2024 (95% [extremely likely] range: 39–65%; Fig. 3a; Supplementary Table 5). Similarly, the combined emissions of the 27 countries of the European Union and United Kingdom contributed to 47% (95% CI: 36–62%), and the emissions of China to 30% (95% CI: 22–41%) of events with over 10% coral bleaching. The emissions of countries that host substantial and iconic warm-water reef ecosystems also made significant contributions to bleaching globally: Australia to 7% (95% CI: 6–9%), Indonesia to 7% (95% CI: 6–8%), and the combined emissions of the 39 countries of the Alliance of Small Island States to 7% (95% CI: 6–8%) of heat-driven events where over 10% of corals bleached. When a more severe bleaching threshold of over 20% bleaching is used then the estimated contribution to the percentage of bleaching events increases; for example, to 54% of events (95% CI: 40–72%) for the emissions of the United States (Extended Data Fig. 8, Supplementary Table 6).

Our approach also enables estimation of a state’s contributions to coral bleaching located within their own territorial waters; for example, China contributed to 37% of events (95% CI: 10–100%) of more than 10% bleaching that occurred within its own territory, the United States to 28% of events (95% CI: 4–56%) within its own territory; Indonesia to 3% (95% CI: 1–7%), and Australia to 4% (95% CI: 0–16%). For China, the United States, and Indonesia, the 99% uncertainty range does not include zero, making it virtually certain that each has contributed to heat-driven coral bleaching of the reef sites under their stewardship (Supplementary Table 5).

A limitation of the ‘but-for’ attribution method is that when impacts are nonlinear in relation to emissions, the assessed contributions of individual emitters can sum to less than or more than 100% depending on the timing of their emissions and the shape of the nonlinearity^47,48^. In the case of coral bleaching we find that, but for the emissions of multiple different emitters a given bleaching event would not have occurred, such that removing each of the individual emitters is sufficient to avoid the bleaching event (see Methods, Extended Data Fig. 9). This fact explains why the contributions of specific emitters can add to more than 100% (Fig. 3a).

Without the emissions of the 172 carbon majors considered here, 84% of heat-driven events with more than 10% coral bleaching over the period 1985–2024 would not have occurred (95% CI: 69–96%; Fig. 3c). Chevron, Saudi Aramco, ExxonMobil and Gazprom are each estimated to have contributed to 14% of events (95% CI: 11–19%) with over 10% of corals bleaching, with BP having contributed to 12% (95% CI: 9–15%) and the National Iranian Oil Company to 11% (95% CI: 9–14%) of bleaching events (Fig. 3c; Supplementary Table 5). The 99% uncertainty range on contribution to bleaching does not include zero for any of these carbon majors, making it ‘virtually certain’ by IPCC standards that each of them has contributed to heat-driven coral bleaching. When a more severe bleaching threshold of more than 20% bleaching is used then the estimated responsibility remains virtually certain and the percentage of bleaching events to which carbon majors contributed increases; for example, to 91% of events (95% CI: 76–100%) for the emissions of all carbon majors and approximately 16% for each of Chevron, Saudi Aramco, and Gazprom (Extended Data Fig. 8, Supplementary Table 6). These results reveal the material contribution individual carbon majors have made in causing coral reef bleaching with contributions to bleaching that exceed those of many states, such as many small island states with high dependence on coral reefs for local livelihoods.

Although a cumulative approach captures the impacts from a specific emitter over several years, extreme climate and weather events have been the focus of much of climate attribution and liability claims^25,33^. The 2023–2024 El Niño led to the most extensive global coral bleaching event so far^49^. Individual state and carbon major contributions to bleaching during this event are virtually certain, but of smaller magnitudes than their contributions to bleaching events over the entire 1985–2024 period (Fig. 3b,d). For example, the United States is estimated to have contributed to 32% (95% CI: 15-49%) of events of more than 10% coral bleaching during 2023–2024 while Saudi Aramco, Chevron, and Exxon Mobil contributed to 6–7% of events (Supplementary Table 7).

To better understand this pattern of specific emitter contributions during 2023–2024 bleaching being smaller than for 1985–2024, we estimated the contributions of specific emitters to each successive global bleaching event. We find that despite large increases due to human-caused climate change in the extent of coral bleaching globally over successive El Niño events from 1998 to 2024, the contributions of individual emission sources such as the United States or China remained relatively constant between the 2015–16 and 2023–24 El Niño events, and contributions from smaller sources like the AOSIS group or Saudi Aramco even declined slightly between these events (Extended Data Fig. 10). This may seem counterintuitive; however, analysis with a conceptual model indicates that this reflects an emergent feature of the application of but-for attribution approaches for impacts which saturate at certain global warming levels (Extended Data Fig. 9)—as is expected to occur for global coral heat exposure and bleaching between 1.5 °C and 2 °C global warming^1,5^. This is because as global warming approaches levels at which bleaching impacts saturate—noting 2024 reached a global warming level of 1.55 °C^50^—the marginal reductions in global temperatures in counterfactuals that remove individual emission sources correspond to an increasingly minor change in impacts. Once the point of impact saturation is passed, contributions from individual emission sources will become indistinguishable within a but-for framework.

### Socioeconomic inequalities in sources and recipients of coral bleaching impacts

Beyond individual states, our approach allows us to quantify the contributions of groups of states to bleaching, such as for those groupings more or less socioeconomically dependent on coral reefs (Fig. 4). To understand the relative contributions of these groups, we use the ‘but for’ approach to assess the contributions from the emissions of each group to coral reef bleaching in the territorial waters of each other group (Methods). Because these contributions do not initially add to 100% due to underlying nonlinearities, we follow previous work^48^ in normalizing the contributions of each group and display them as proportions of the total contribution to bleaching (Methods).

**Fig. 4.**
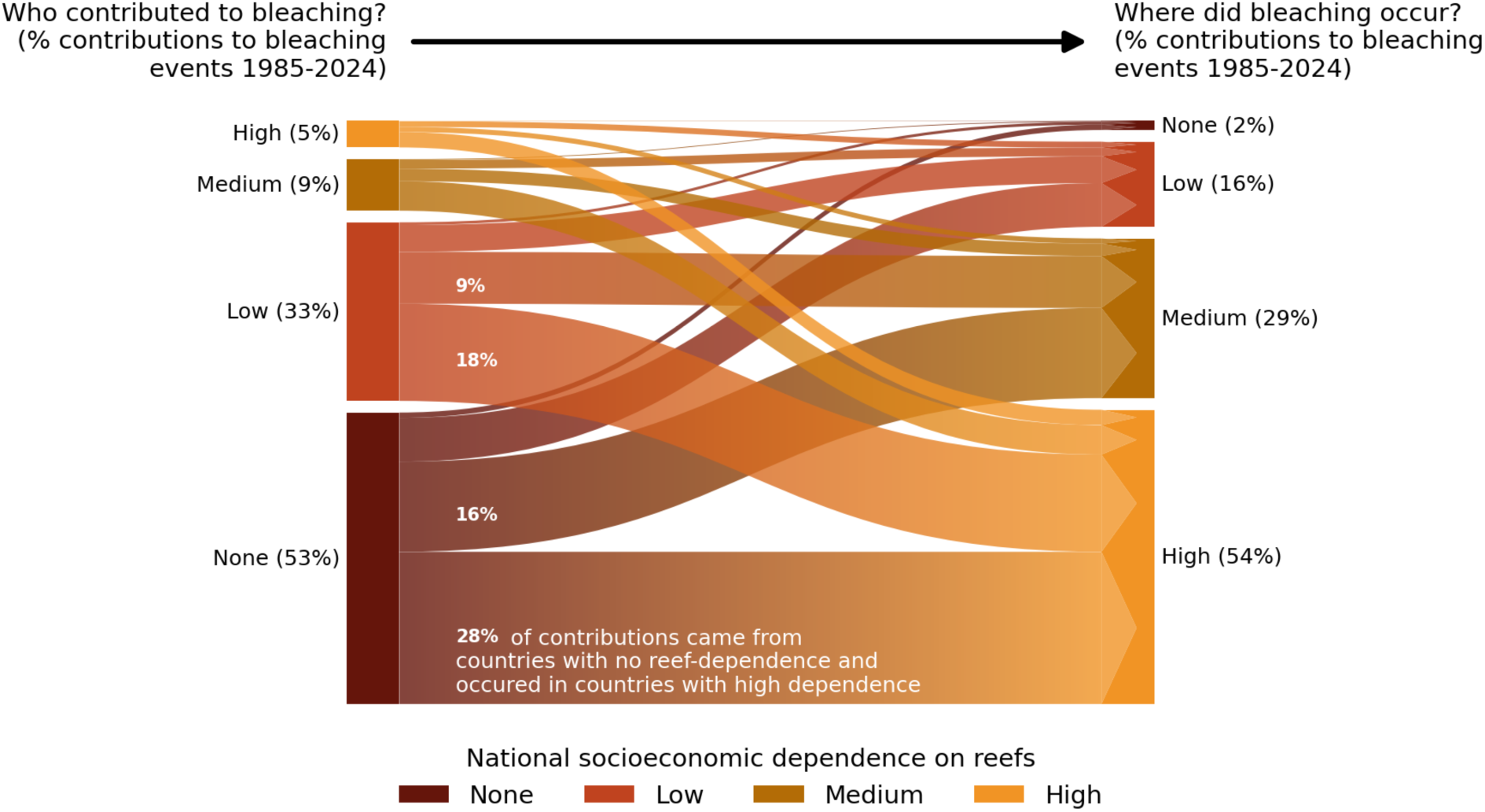
States contributing most to coral bleaching have no socioeconomic dependence on reefs. Arrows indicate flows from sources of bleaching (left) to recipients of bleaching (right). Left-hand bars indicate the percentage of total contributions to bleaching events of ≥ 10% of corals across 1985–2024 which came from groups of states with differing socioeconomic dependence on reefs. Right-hand bars indicate the groups within which these bleaching events occurred. Colours indicate levels of socioeconomic dependence on reefs based on reef-associated population, reef fisheries employment, reef-associated exports, nutritional dependence on fish and seafood, reef-associated tourism, and shoreline protection with data taken from Resource Watch^9^.

There is a dual inequality to the heat-driven bleaching of coral reefs: States with no or low socioeconomic dependence on coral reefs have historically contributed the most to heat-driven bleaching, but this bleaching has occurred primarily in states with high socioeconomic dependence on reefs (Fig. 4). Fifty-three percent of all contributions to bleaching events came from states with no socioeconomic dependence on reefs—including the United States, the United Kingdom, and Russia—with most of this contribution causing bleaching in states with high socioeconomic dependence on reefs (e.g., Antigua and Barbuda, Vanuatu, Philippines). By contrast, the emissions of the group of highly reef-dependent states only generated 5% of total contributions to bleaching. This pattern holds across major El Niño events, for different bleaching thresholds, and for patterns of DHW exposure rather than bleaching (Supplementary Figs. 4–7). Together, these results emphasise that the negative human impacts of coral reef bleaching in terms of lost ecosystem services and livelihoods have been felt by communities who are far removed and have not benefited from the greenhouse gas emissions which have predominantly caused bleaching.

## Discussion

While the mechanistic role of high sea water temperatures in causing coral bleaching has long been known^13^, our models formally quantify in an end-to-end attribution the extent of bleaching attributable to anthropogenic greenhouse gas emissions, with stark conclusions: in the absence of human-caused climate change, virtually none of the observed heat-driven bleaching across 1985–2024 would have occurred, even during strong El Niño events (Fig. 2). This finding suggests warm-water corals were well-adapted to pre-industrial climate conditions (defined here as 1850–1900), able to experience major El Niño events with minimal bleaching, and underlines the overwhelming role of human-caused climate change, as opposed to natural climate variability, in causing recent, massive coral reef damage worldwide.

This massive extent of human-caused climate change’s impact on corals is assessed in this study based on coral thermal stress and bleaching, but this is likely a conservative assessment of the full contribution to impacts on corals from anthropogenic greenhouse gas emissions because we do not assess the influence of emissions on ocean acidification, on other climate hazards such as sea level rise and tropical storm intensification, or on changes in El Niño intensity or frequency—all of which can impact corals^51,52^. Nevertheless, change detection for tropical storms and El Niño cycles remains difficult at current levels of global warming^53–55^ and the impacts of heat on corals are likely to outweigh those of acidification over the 21st century^51^, justifying the current focus on bleaching from ocean warming.

We also quantify contributions to heat-driven coral bleaching by specific states and carbon majors (Fig. 3). It is virtually certain for many individual countries and carbon majors that their emissions have caused coral bleaching, with the largest carbon majors having each contributed to 14% of events with more than 10% coral bleaching. In contrast to biodiversity impacts from land-use and sea-use change—whose upstream drivers are difficult to trace across globally-distributed networks of trade and extractive industries^56^—the emissions of states and carbon majors are relatively well documented^57,58^, enabling assessment of their responsibility for climate-driven impacts. Applying end-to-end attribution to quantify contributions of specific emitters to climate change impacts for a wider diversity of ecological communities and processes contributing to people via food, water, and other benefits (e.g., pollination, pest regulation) could support efforts to hold specific emitters liable for damages caused by climate warming^59^. For example, an ongoing case brought by Indonesian islanders against Swiss cement company the Holcim group cites the impact of Holcim’s emissions on their island via both rising sea levels and from coral reef damage due to warmer sea temperatures^60^. Our framework estimates that Holcim’s emissions contributed to approximately 3% of observed coral bleaching events where over 10% of coral colonies bleached across the Indonesian sites in our sample (Supplementary Table 5).

The extent to which attribution science on source-specific contributions finds use in the courts relating to human and interspecies justice depends in part on jurisdictional contexts and discretion. Nevertheless, our results shed light on a new potential challenge in this area: applying but-for attribution to impacts saturating rather than increasing with higher levels of global warming returns declining contributions of individual emitters when examining time periods nearing and passing the point of saturation. We show that this is the case for individual emitter contributions to coral bleaching using time periods of successive major El Niño events as Earth begins to reach the 1.5 °C global warming level projected to lead to loss of 70–90% of warm-water coral reefs^1^ (Extended Data Figs 9 and 10). To be clear, the use of but-for attribution in this context can still show an emitter is more likely than not, or even virtually certain (e.g., when confidence intervals for contribution do not overlap zero), to have caused a damage; however, any use of estimated contributions for allocation of damages among emitters may become more complicated. Ways to account for these dynamics considering jurisdictional precedent will be important not just for corals but also a range of other systems exhibiting climate-driven saturating impacts or tipping points.

Strong inequalities exist among the sources and recipients of coral bleaching impacts with states that contributed most to coral bleaching having the least socioeconomic dependence on coral reefs (Fig. 4). This underlines the continued importance of integrating climate justice into the resourcing and implementation of climate mitigation, adaptation, and loss and damage mechanisms in the context of coral-reef-dependent communities. As global warming surpasses 1.5°C—a level understood to be catastrophic for warm-water coral reefs—the extent to which natural and assisted adaptation might enable reef persistence in some locations over the second half of the twenty-first century remains highly uncertain^51,61^. By assessing the role of anthropogenic greenhouse gas emissions, and of specific emitters, in causing coral bleaching our results make an important contribution to establishing accountability for non-economic loss and damage from climate change.

## Methods

### Coral bleaching data

Coral bleaching observations were drawn from two sources. The first is a recently published database, the Global Coral Bleaching Database^3^, that spans 1980–2020 and is one the most complete, publicly-available databases on the extent of coral bleaching at 14,405 sites across 93 countries compiled from seven data sources, including those collated by Donner *et al*.^62^ and the terminated ReefBase database^63^. To include more recent data we also made use of the publicly available subset of the MERMAID^28^ (Marine Ecological Research Management Aid) database, which provided an additional 860 observations from 564 sites spanning 1998–2024. MERMAID survey records were cross-referenced against the Global Coral Bleaching Database by site coordinates, survey year, and bleaching percentage to identify potential duplicates. No overlapping records were found, consistent with the fact that the Global Coral Bleaching Database predates the MERMAID database.

Data from the Global Coral Bleaching Database^3^ includes coral population-level and colony-level data for a site. Following previous studies^39,64^, we make use only of population-level data on the percentage of coral colonies that exhibited bleaching to ensure our outcome variable reflects the extent of site-wide bleaching rather than the intensity of bleaching across an unknown portion of coral colonies at a site. Retaining only population-level bleaching records from the Global Coral Bleaching Database and MERMAID reduced our temporal coverage to the period 1998–2024. Population-level bleaching extent was reported either as a direct site-level percentage or as per four transect estimates for a site, in which case the mean across the four transects was computed. This resulted in a unified continuous response variable for coral bleaching, ranging from 0–100%. Each site was assigned to an ecoregion by spatially intersecting site coordinates with the Corals of the World ecoregion shapefiles^65^, following the approach used by the Global Coral Bleaching Database^3^. When joined, the two datasets produced a combined 10,897 population-level observations across 4,479 unique sites and 86 ecoregions for the 1998–2024 period.

### SST data and degree heating weeks calculation

To quantify exposure of coral reef sites to heat stress we follow decades of coral reef science and monitoring by using Degree Heating Weeks (DHW)^12,29,66^. We derive DHWs from daily, global 5km sea-surface temperature (SST) data from NOAA Coral Reef Watch, a high-resolution product available from 1985 to near real-time that is derived from multiple reanalyses and satellites specifically for use in high-resolution monitoring and alerts of coral bleaching^29^. We use the “hotspot” variable^29^, *T_x_*_,*t*_, reflecting daily anomalies of SSTs from 1**°**C over the maximum monthly mean (MMM) temperature derived over the period 1985-2012, to calculate the DHW metric:

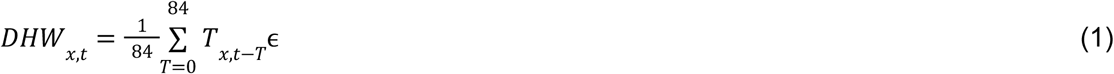

Where ɛ = 0, 1 depending on whether the daily SST exceeds 1°C over MMM. This metric reflects cumulative heat exposure over the prior 12 weeks (84 days, approximately 3 months) and is widely used for bleaching alerts and to plan monitoring of coral bleaching. We use the maximum monthly value of daily DHWs to reflect peak exposure within a month, matching these with bleaching observations for further analysis.

We calculated additional SST-derived variables to reflect historical coral heat exposure variability and trends for use in sensitivity tests in our bleaching impact model. These were: the rate of historical SST warming, defined as the difference in daily SST anomalies, *T_x_*_,*t*_ , averaged over the period 1985–2004 and 2005–2024; the historical standard deviation of SST anomalies over the period 1985–2004; and the historical standard deviation of DHW over the period 1985–2004.

Data on the global mean surface temperature (GMST) is calculated from ERA5 re-analysis^67^ surface temperatures, taking area-weighted averages and averaging across days of a year.

### Statistical coral bleaching impact model

Modelling of the relationship between marine heat exposure and coral bleaching has typically followed one of two approaches: (i) modelling changes in the percentage of bleaching of coral communities^39,64^, or (ii) modelling the probability of bleaching after making bleaching percentages into a binary variable based on a given bleaching threshold^42,68–70^. Here, we follow the first approach by modeling changes in the percentage of bleaching. We do this so that marginal changes in bleaching can be fully resolved thereby enabling the evaluation of the contribution to coral bleaching of marginal changes in anthropogenic emissions (see Methods on “End-to-end impact attribution”).

To identify the plausibly causal effect on coral bleaching of changes in DHW, we use a fixed-effects modelling framework for causal identification commonly applied in climate econometrics^71,72^, and increasingly in climate impacts research in epidemiology^73,74^ and ecology^23,75^. This framework is designed to strengthen a causal interpretation of effects by semi-parametrically accounting for unobserved spatial and temporal confounding factors in order to isolate variation in the climate variable of interest which is highly exogenous and unlikely to be correlated with other socio-ecological factors. Causal estimates of the effect of a particular climate hazard on a given outcome enable counterfactual simulations of climate impacts in which the climate is changed and all other factors are held constant, thereby enabling end-to-end attribution in the context of climate change^33^ , as is the aim in our study.

We use a standard approach of a two-way fixed effects panel regression model and for this analysis we restrict the dataset to include only sites with repeated observations, yielding a dataset of 6,986 observations across 1,626 sites and 60 ecoregions globally for 1998–2024 (Figure 1a). In the panel regression we use fixed-effects by site and year^72^ to control for unobserved site-level confounders (e.g., differences in baseline levels of bleaching at particular sites due to depth or distance from shore), and for unobserved confounders in a given year that are common across sites (e.g., the strong influence of an El Niño event). These approaches fully remove confounders by nonparametrically de-meaning the data along these dimensions and thereby provide a stronger basis for causal inference compared to approaches with random-effects models^39,64^ where confounders are only partially removed with random intercepts. We furthermore include a fixed-effect for the interaction between ecoregion and month to account for seasonality of heat exposure and bleaching that is common within ecoregions.

Our baseline panel regression model is a linear fixed-effect model, in which we allow the effect of DHW on bleaching to vary based on absolute latitude. Let *y_it_* ɛ [0, 1] denote the bleaching percentage for site *i* at time *t*. The model is specified as:

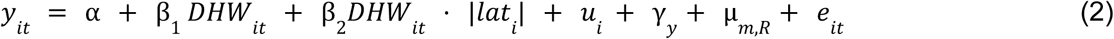

where α is the global intercept, β_1_ is the effect of DHW exposure, β_2_ captures the modulation of thermal sensitivity by absolute latitude, and *ui* and γ*y* are site and year fixed effects respectively, equivalent to allowing each site and year to have its own intercept such that site-level fixed effects absorb unobserved and time-invariant variation in baseline bleaching across sites (e.g., due to depth) and year fixed-effects absorb confounders common across sites in a given year (e.g., El Niño events). We note that controlling for El Niño in this step ensures that the bleaching-exposure relationship which we model is not spuriously driven by other El Niño mediated phenomenon, while in the later counterfactual analysis we purposefully retain the timing of El Niño events given their major role in driving DHW exposure. The fixed effect µ*_m_*_,*R*_ denotes a month, *m*, by ecoregion, *R*, fixed-effect that accounts for ecoregion-specific seasonality in bleaching that may spuriously relate to seasonally varying SSTs.

We evaluate the linear model specification by comparison to a non-parametric binned fixed-effects model which flexibly tests for potential non-linearities in the bleaching response to DHW, and by evaluating the residuals as a function of DHW and percentage bleaching. We find the linear model is largely consistent with the response of bleaching to DHW recovered by the non-parametric, binned approach (Fig. 1b) and that residuals of the linear model remain unbiased across the majority of the DHW distribution up to 20% bleaching, implying no systematic misspecification by the choice of a linear fit (Supplementary Figure 2). We also test higher-order polynomials of DHW (quadratic and cubic) and show these provide no additional predictive power (Supplementary Table 2), consistent with the non-parametric, binned model estimates of the bleaching response revealing limited additional non-linearities compared to the linear model (Fig. 1b). We further test the linear specification in comparison to a zero-one inflated beta model designed specifically for zero-inflated data (as is the case with coral reef bleaching observations), finding very consistent results (Supplementary Figure 1). We therefore choose to retain the linear fixed-effects model as our main specification because the zero-one inflated beta model requires random rather than fixed-effects which provide a less clear removal of confounders and therefore a weaker causal interpretation^76^.

We analyse the spatial autocorrelation of the linear model residuals and find that for sites within the same ecoregion residuals are strongly correlated below distances of 200km but very weakly correlated at greater distances (Supplementary Figure 3). We therefore evaluate the uncertainty of our main specification using errors as specified by Conley^77^, assuming non-independence between sites separated by less than 200km. The effect of DHW on bleaching remains strongly significant when instead clustering by larger ecoregions (Supplementary Table 4).

We conduct further sensitivity tests on the estimation of DHW values in the linear fixed-effect model. First, we evaluate the relationship between DHW and bleaching at different lead and lag times, finding that the magnitudes and significance of the relationship between DHW and coral bleaching decay to insignificant values at lags or leads greater than two months (Supplementary Table 9). This is consistent with the definition of Degree Heating Weeks as cumulative exposure over three months. We also test the robustness of the impact model to alternative definitions of DHW accumulated over 11 weeks and based on exceedances of 0.4 °C above maximum monthly climatology, as advocated by Whittaker et al.^44^ and find very similar results to DHW defined as the cumulative 12-week sum of degrees of SST 1°C above the maximum monthly mean SST climatology as used in global coral bleaching alert systems^29^ (Supplementary Table 3).

We explore potential heterogeneities in the response of bleaching to DHW exposure based on multiple climate, ecological, or social factors identified in prior literature^39–42,51^ (Supplementary Table 1). We test for moderation of the effect of DHW on bleaching by depth (data taken from Global Coral Bleaching Database) and absolute latitude. To capture the potential for adaptation based on exposure to historical temperature variability or warming to potentially reduce bleaching sensitivity^39,42^, we test for a moderating effect of historical variability in SSTs and DHW (calculation of SST warming rates and variability are described in the SST data section above). We also test whether local human pressures moderate heat-driven bleaching sensitivity using mean turbidity, an indicator of water quality, and a site’s Marine Protected Area status (data taken from Global Coral Bleaching Database).

Lastly, we note there is relatively limited observational data of population-level bleaching above 20% in the global datasets used in this study. As such, the fact that model residuals appear to exhibit some more systematic biases beyond a 20% level of bleaching (Supplementary Figure 2c,d), means we limit our use and interpretation of the bleaching impact model to present end-to-end impact attribution results based on the number of coral reef sites and bleaching events with greater than 10% population-level bleaching or greater than 20% population-level bleaching. Ten percent population-level bleaching is widely used as a threshold level for moderate bleaching within coral reef surveys and research^2,11^.

### End-to-end impact attribution

We build on and extend existing end-to-end impact attribution workflows^33^ to develop a workflow for coral bleaching which traces the impacts of heat on coral bleaching back to the emissions of states and carbon majors which contributed to global warming. This workflow is based on the following steps:

1. Generating counterfactual scenarios of global mean surface temperature (GMST) in the absence of the emissions of individual companies and countries;
2. Deriving pattern-scaling relationships between GMST and SSTs at coral reef sites from observed historical data;
3. Generating counterfactual scenarios of coral heat exposure from steps (1) and (2);
4. Applying the coral bleaching impact model to the counterfactual coral heat exposure scenarios from step (3) to generate counterfactual estimates of coral reef bleaching, while propagating uncertainties across all steps.

In the following sections we provide more details on the methodology of each of these steps.

### Global temperature counterfactuals

We use the Finite Amplitude Impulse Response (FaIR) simple climate model to generate counterfactuals of GMST in the absence of anthropogenic climate forcing from various emission sources. A “natural” simulation with only natural forcing provides an estimate of GMST in the absence of all anthropogenic emissions from fossil fuel use, land-use, and land-use change. We furthermore conduct “leave-one-out” simulations in which we remove the emissions of 172 major fossil-fuel companies (considering their scope 1 and 3 emissions of both CO_2_ and methane with varying temporal coverage between 1850-2024 from the carbon major database^58^) and from countries or groups of countries (using national territorial emissions of CO_2_ from 1850-2024 from the global carbon budget^57^) following the methods of ref.^33^. Two country blocs are included for territorial emissions: the 27 countries of the European Union and the UK, and the 39 countries of the Alliance of Small Island States. This design for generating counterfactuals follows the “but for” logic applied in legal contexts to assign responsibility to actors contributing to an event, by evaluating the outcome which would have occurred *but for* the actions of that party. We also grouped emissions of individual countries by their levels of socioeconomic dependence on coral reefs based on data from *Resource Watch*^78^. These data reflect a range of benefits provided to people by coral reefs, evaluated based on reef-associated population, reef fisheries employment, reef-associated exports, nutritional dependence on fish and seafood, reef-associated tourism, and shoreline protection. For simplicity, we harmonize the two upper levels of high and very high dependence into a single category, and in cases where parts of a country are assigned different levels of reef dependence (e.g. Florida vs the rest of the USA), we use a population weighting to assign an overall category of reef dependence to that country.

We use version 2.2.4 of FaIR to generate counterfactual scenarios removing the emissions of each individual company, country, and group of countries. To account for uncertainty in the transient climate sensitivity to emissions, we sample the full set of 841 posterior ensemble members from calibration version 1.4.1 of FaIR, which was based on a Bayesian framework to evaluate FaIR simulations in relation to CMIP6 simulations and observed climate changes^32^, resulting in 841 counterfactual scenarios for each company, country or group of countries. This recent calibration includes volcanic and solar forcing with which we are able to simulate global temperature trajectories from 1750–2024. The trajectories of GMST under the various counterfactuals of interest are shown in Supplementary Figure 10.

### GMST to local SST pattern scaling

For each calendar day, *d*, we derive a scaling relationship between the observed global mean surface temperature (*GMST*_y_) and observed SST anomalies, *T_x_*_,*t*_ , at each coral site *x*:

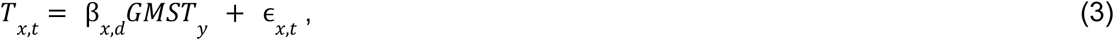

With error ɛ*x*_,*t*_ . The statistics of the deviations of the resulting scaling relationships, β*_x_*_,*d*_ , within seasons and between calendar days are shown in Supplementary Figure 11. Given that differences in the scaling between adjacent calendar days are very small, we continue without pooling data across adjacent calendar days in order to preserve the specificity of our estimates across days, while fully accounting for uncertainty in these scaling parameters using block-bootstrapping (described below). We use observational data, rather than simulations from global climate models, to derive scaling relationships in order to benefit from the high spatial resolution (5km) SST data used for measuring historical exposure of coral reefs to heat. We use a linear relationship following work on attribution of daily SSTs to climate change^35^ and the fact that local temperature extreme intensity scales linearly with global temperatures^79^. We then use a recently developed block-bootstrapping framework to capture the uncertainty in these pattern-scaling relationships^34^. Specifically, we evaluate the autocorrelation-function of the residuals of equation (2) and their significance according to Bartlett’s formula, finding a very weak autocorrelation structure which decays to negligible and insignificant values after a single year (Supplementary Figures 12 and 13). As such we proceed with 2-year block-bootstraps of equation (2), to estimate a distribution of β*_x_*_,*d*_ , and take 841 samples with replacement to match the number of uncertainty samples stemming from the GMST counterfactuals produced by FaIR. We note that this procedure has been shown to reproduce the levels of uncertainty derived from large ensemble climate model simulations for temperature extremes, and can therefore be expected to accurately capture uncertainty from internal climate variability relatively well^34^. This allows us to derive robust uncertainty estimates for these scaling parameters while exploiting the high resolution observed SST data. Our approach therefore avoids the need to use global climate model data which would require bias-adjustment to reflect SSTs at local coral sites and would also risk inclusion of significantly biased SST trends from climate models compared to observations in important ocean regions for coral reefs, such as the tropical Pacific^37^.

### Counterfactual coral heat exposure

We combine the counterfactual scenarios of GMST 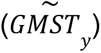 with the scaling relationships between historical GMST 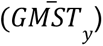 and observed SST anomalies (*Tx*,*y*,*d*) to generate daily counterfactuals of SST 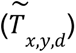 following the formula:

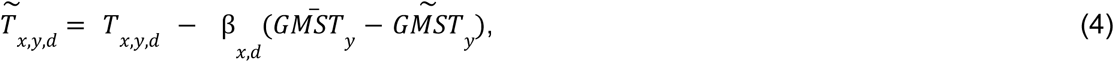

Noting that 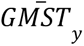 is the GMST simulated by FaIR in the historical forcing scenario (that is, with no anthropogenic emissions removed) and 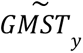 is the GMST simulated by FaIR in a scenario with some amount of anthropogenic emissions removed (e.g., those from a specific country or company). We generate counterfactual SST anomalies for each day for the historical period 1985-2024. For the 366th day of the year in leap-years, we use the value of the scaling relationship β*_x_*_,*d*_ from the adjacent 365th day of the year. Using these counterfactual SST anomalies, we then re-calculate Degree Heating Week values for each coral reef site using equation 1 (examples shown in Supplementary Figure 14).

### Counterfactual coral bleaching and uncertainty propagation

In the final stage of the end-to-end attribution framework, we use the coral bleaching impact model to quantify coral bleaching under different scenarios of DHW exposure including observed and counterfactual DHWs (that is, with the emissions of individual companies or countries removed). We do this by applying the coefficients β_1_ and β_2_ from the statistical coral bleaching impact model in equation (2) to the historical and counterfactual DHW time series of a given coral site to generate estimates of coral bleaching with and without a given source of anthropogenic emissions. Here, we use the full set of 4,479 known coral sites from the combined Global Coral Bleaching and MERMAID database. This allows us to provide a more spatially comprehensive assessment and is justified by the fact that SST observations are available across all known coral sites and that overall statistics of anthropogenic contributions to exposure and bleaching are very similar when evaluated across the smaller set of sites used in the panel regression bleaching–DWH impact model (see robustness tests shown in Supplementary Figs 8 and 9).

Across the end-to-end attribution framework, three important sources of uncertainty exist, reflecting:

i. uncertainty in the transient climate sensitivity to emissions;
ii. uncertainty in the pattern scaling relationship between GMST and local SSTs, driven by different realisations of regional warming due to internal climate variability;
iii. statistical uncertainty in the relationship between heat exposure (measured as DHW) and coral reef bleaching.

Samples from each of these three sources of uncertainty are independent and thus we combine them to capture the full range of climate and statistical uncertainty in counterfactual estimates of DHW exposure and coral bleaching. First, we account for uncertainty in transient climate sensitivity to emissions by using the full 841 posterior samples available for FaIR model parameterisation: For each counterfactual scenario of GMST, we generate 841 GMST time series, one for each of the 841 posterior samples of the FaIR model. To aid calculation of overall uncertainty, the number of uncertainty samples for climate sensitivity was used to set the sample size for all three sources of uncertainty. Second, we account for uncertainty in the pattern-scaling relationships between GMST and SSTs by taking 841 samples of the block-bootstrap procedure to capture uncertainty in the pattern-scaling coefficients for each coral reef site and calendar day (see Methods “GMST to local SST pattern scaling”). Third, uncertainty in the estimation of coefficients for the coral bleaching impact model are obtained by sampling from the variance-covariance matrix of the impact model coefficients (β_1_ and β_2_ from equation 2) which accounts for the observed levels of spatial autocorrelation up to 200km, according to Conley. Finally, we combined these three sources of uncertainty into the counterfactual–SST–impact model workflow. Uncertainty estimates presented for DHW counterfactual values (e.g., Figure 2a) include uncertainty from only two sources: transient climate sensitivity and pattern scaling GMST to SSTs. Uncertainty estimates of coral bleaching include all three uncertainty sources. We present uncertainties with confidence intervals reflecting the uncertainty language used by the Intergovernmental Panel on Climate Change^16^

### Nonlinearity in “but-for” attribution calculations

Many entities have emitted greenhouse gases and therefore contributed to ecosystem impacts like coral bleaching. It is therefore necessary to design an experiment that isolates the contribution of an individual actor of interest to these impacts. Many such experimental designs have been proposed^48^. In this study, we follow a substantial body of previous work^33,80–83^ in designing a “but-for” modeling experiment: We subtract a single actor’s emissions from global emissions and compare the resulting coral bleaching.

It has long been recognized, however, that this approach can lead to counterintuitive results when the relationship between emissions and impacts is nonlinear^47,84^. In the case of coral bleaching, while our underlying regression model is linear (Fig. 1), there are multiple sources of nonlinearity: we evaluate the results at particular thresholds of bleaching extent (e.g., 10% bleached), and corals can saturate in their level of bleaching (e.g., once all corals die, no further damage can be done). If a given bleaching event would not have occurred without the emissions of multiple different sources, each of those sources are calculated as contributing to the event, and the total contributions can sum to more than 100%.

We illustrate this phenomenon further with a simple conceptual model of attributing climate impacts to individual sources in the presence of nonlinearity, showing that it is possible to achieve total contributions well above 100% with sufficiently nonlinear impact functions (Extended Data Fig. 9). As the field of source attribution develops, further analysis will be necessary to fully understand the varied types of nonlinearity in impact functions and their implications for emitters’ contributions.

## Data availability

Data on the bleaching of coral reefs is publicly available from the Global Coral Reef Bleaching Database (https://springernature.figshare.com/articles/dataset/Global_Coral_Bleaching_Database/17076287?file=31573421) and MERMAID database (https://explore.datamermaid.org/). Gridded SST data used in this analysis is publicly available from the National Oceanoic and Atmospheric Administration (https://coralreefwatch.noaa.gov/product/5km/index_5km_sst.php). The ERA5 reanalysis is publicly available from Copernicus (https://cds.climate.copernicus.eu/datasets/reanalysis-era5-single-levels?tab=overview). Data on national and company level emissions are respectively publicly available from the Global Carbon Budget (https://globalcarbonbudget.org/) and Carbon Majors Database (https://carbonmajors.org/). Data on socioeconomic dependence of coral reefs are publicly available from Resource Watch (https://resourcewatch.org/dashboards/coral-reefs).

## Code availability

Code for the reproduction of the analysis is available from two repositories, one dealing with the generation of DHW counterfactuals (https://github.com/maxkotz17/Coracle_maxkotz) and one dealing with the DHW-bleaching impact model and its application to the DHW counterfactuals (https://github.com/climaterisklab/Coracle).

## Acknowledgements

We thank scientists and citizen science contributors to the global monitoring of coral bleaching.

## Author contributions

C.H.T., P.P., and M.K. designed the research; M.K. and P.P. performed analyses; R.G., O.B., M.D., C.C., A.S.M. provided feedback on initial results and refinement of methods; M.K., P.P., A.S.M. and C.H.T developed figures; M.K. and C.H.T wrote the first draft of the manuscript; All authors contributed to writing the final draft.

## Extended Data Figures

**Extended Data Figure 1.**
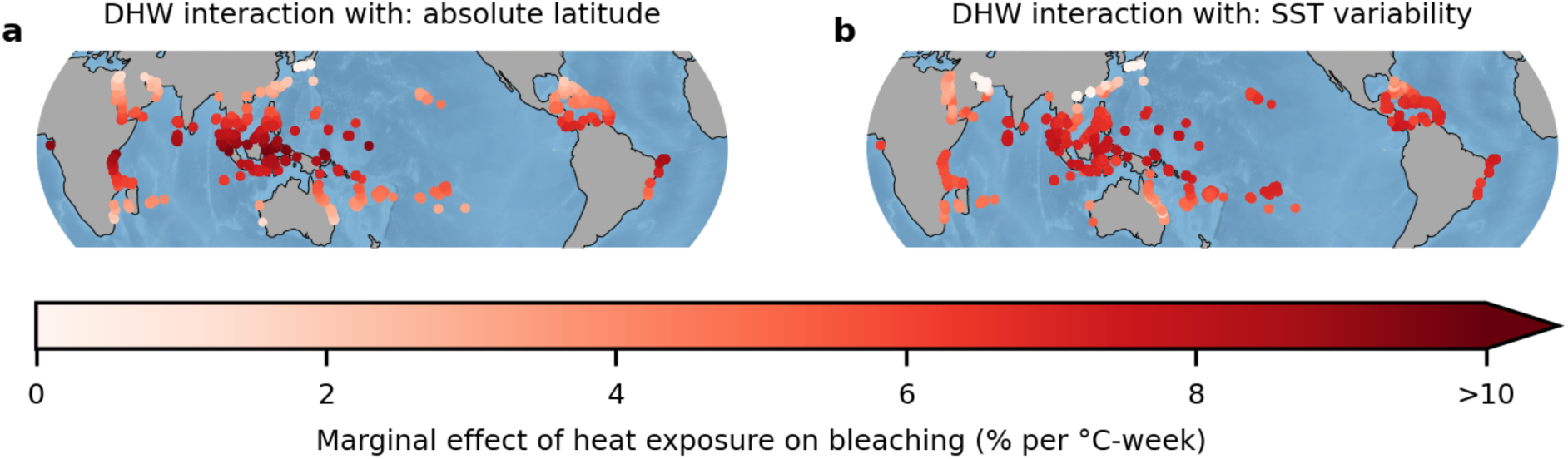
Heterogeneity in bleaching sensitivity to heat exposure accounting for moderating effects of latitude and SST variability. (a) Marginal effect of heat exposure on bleaching when including an interaction with latitude or (b) interaction with historical SST variability (measured as the standard deviation of monthly values over 1985-2005).

**Extended Data Figure 2.**
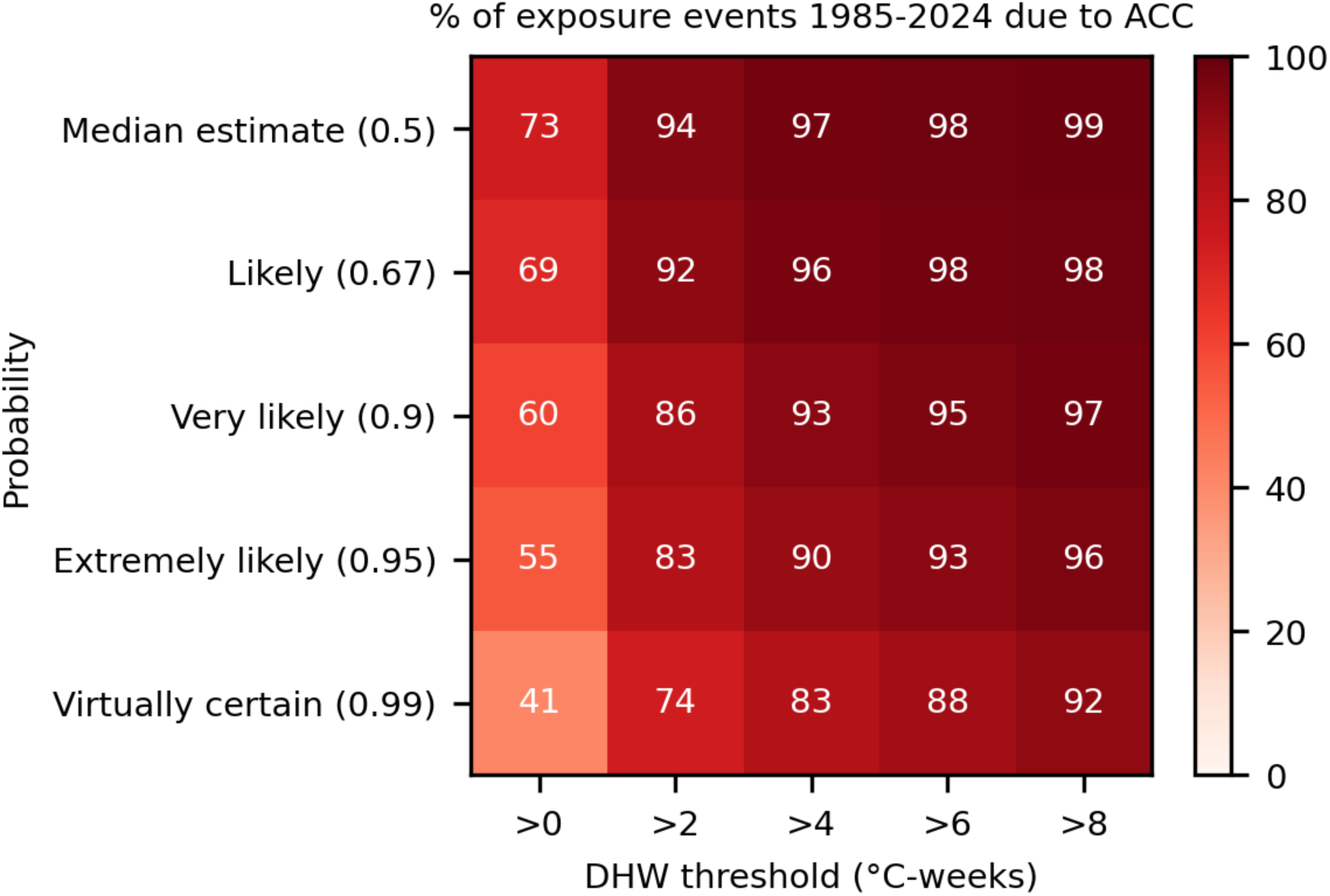
The percentage of heat exposure events for coral reefs due to anthropogenic climate change at different DHW thresholds and likelihoods. Colours and numbers indicate, for a given DHW threshold, the percentage of heat exposure events in the observed climate which did not occur in the counterfactual climates with all anthropogenic climate forcing removed. Estimates for the median value across uncertainty samples and at different quartiles corresponding to IPCC likelihoods (the 67th, 90th, 95th and 99th percentile across uncertainty samples) are given. These probabilities represent a one-sided likelihood, rather than the upper range of a confidence interval, and should be interpreted as: “virtually certain (99% chance) that at least 83% of DHW exposure events of 4 °C-weeks were due to anthropogenic climate change”.

**Extended Data Figure 3.**
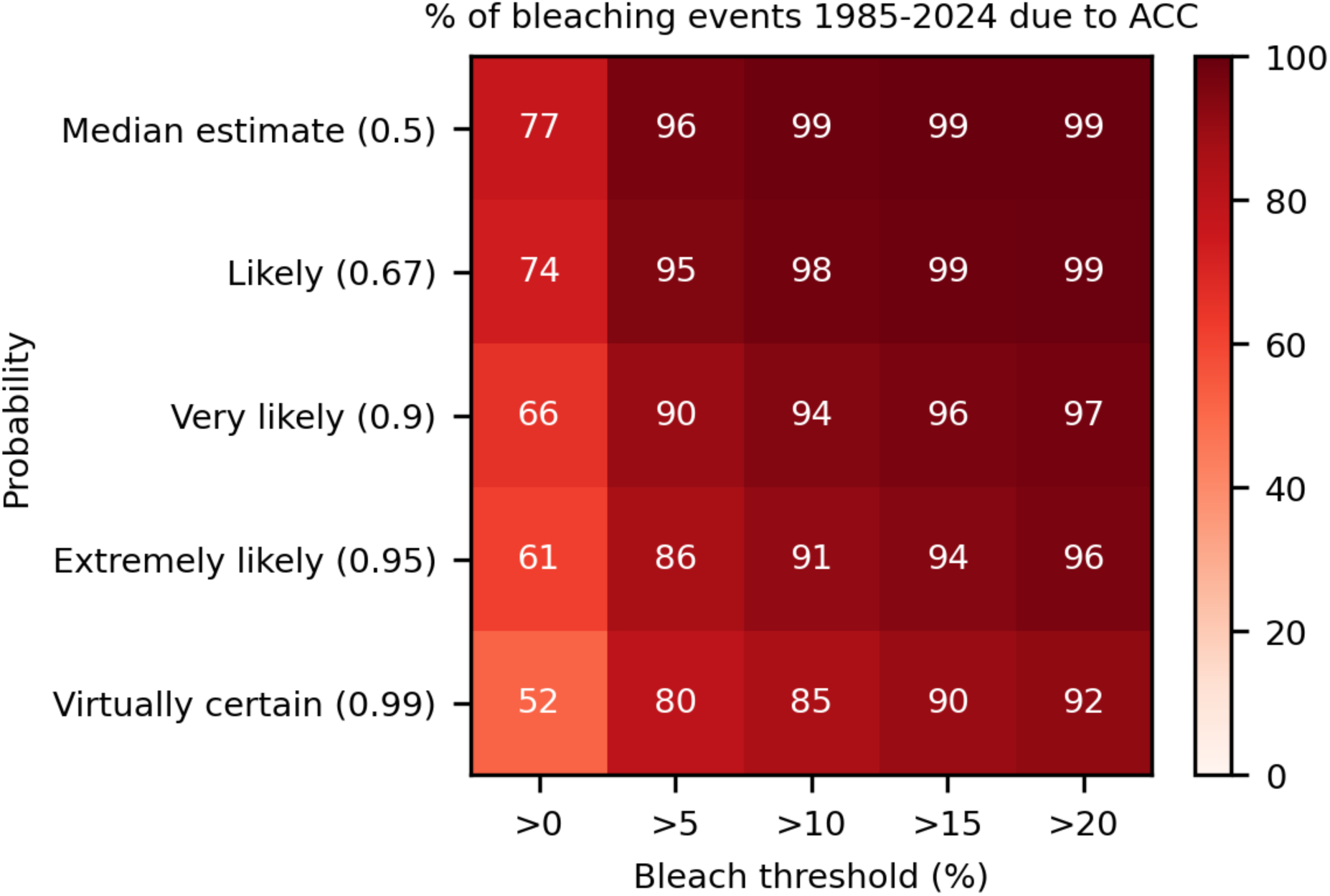
The percentage of coral bleaching events due to anthropogenic climate change at different bleaching thresholds and likelihoods. Colours and numbers indicate, for a given bleaching threshold, the percentage of bleaching events in the observed climate which did not occur in the counterfactual climates with all anthropogenic climate forcing removed. Estimates for the median value across uncertainty samples and the values for different likelihood terms corresponding to IPCC uncertainty guidance (the 67th, 90th, 95th and 99th percentile across uncertainty samples) are given. These probabilities represent a one-sided likelihood, rather than the upper range of a confidence interval, and should be interpreted as: “virtually certain (99% chance) that at least 85% of 10% bleaching events were due to anthropogenic climate change”.

**Extended Data Figure 4.**
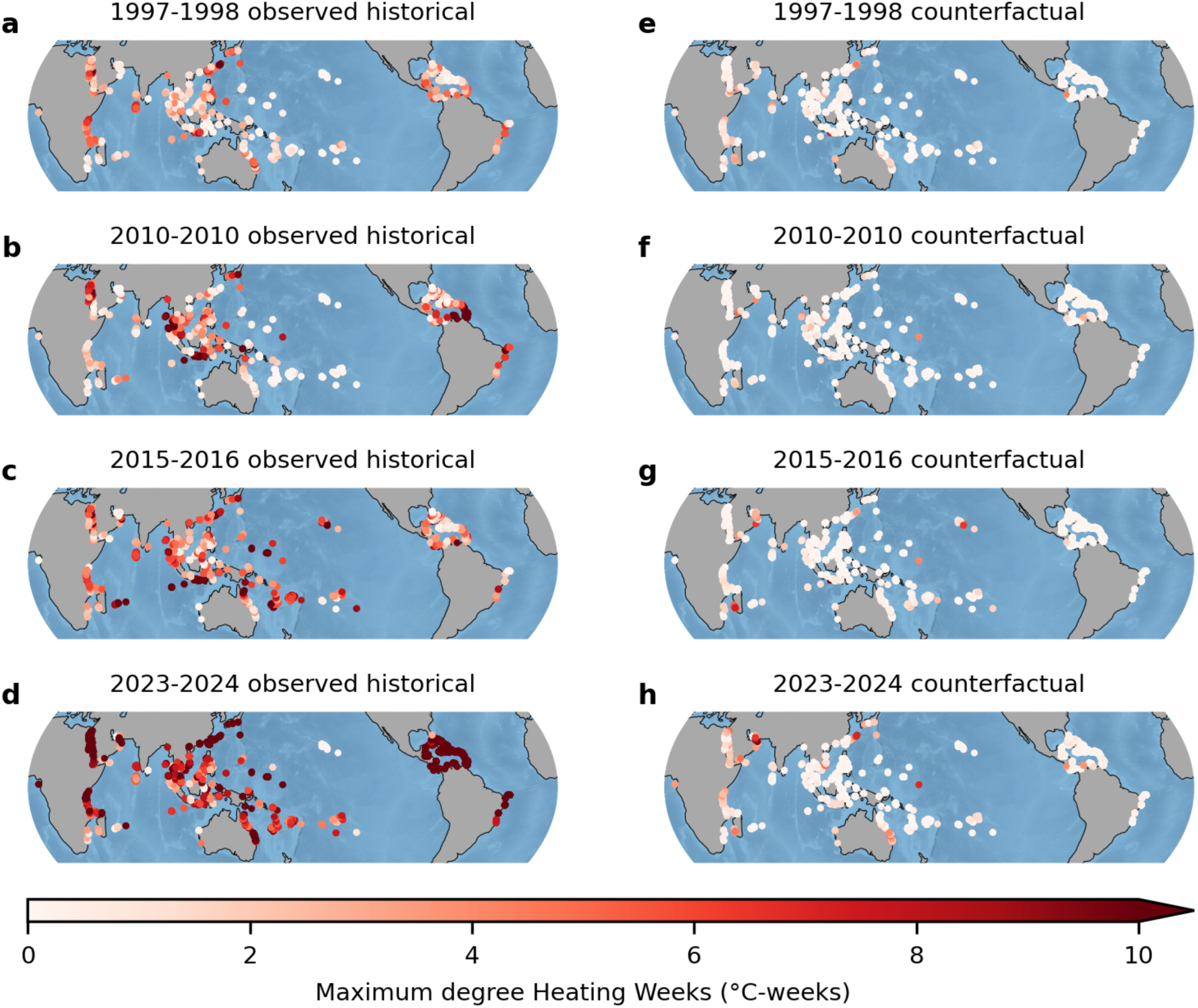
Degree heating week exposure at coral reef sites for major El Nino events is higher in the observed compared to counterfactual climate. (a-d) Maximum degree heating weeks during major El Niño events as factually observed, and (e-h) under counterfactual climates with all anthropogenic emissions removed.

**Extended Data Figure 5.**
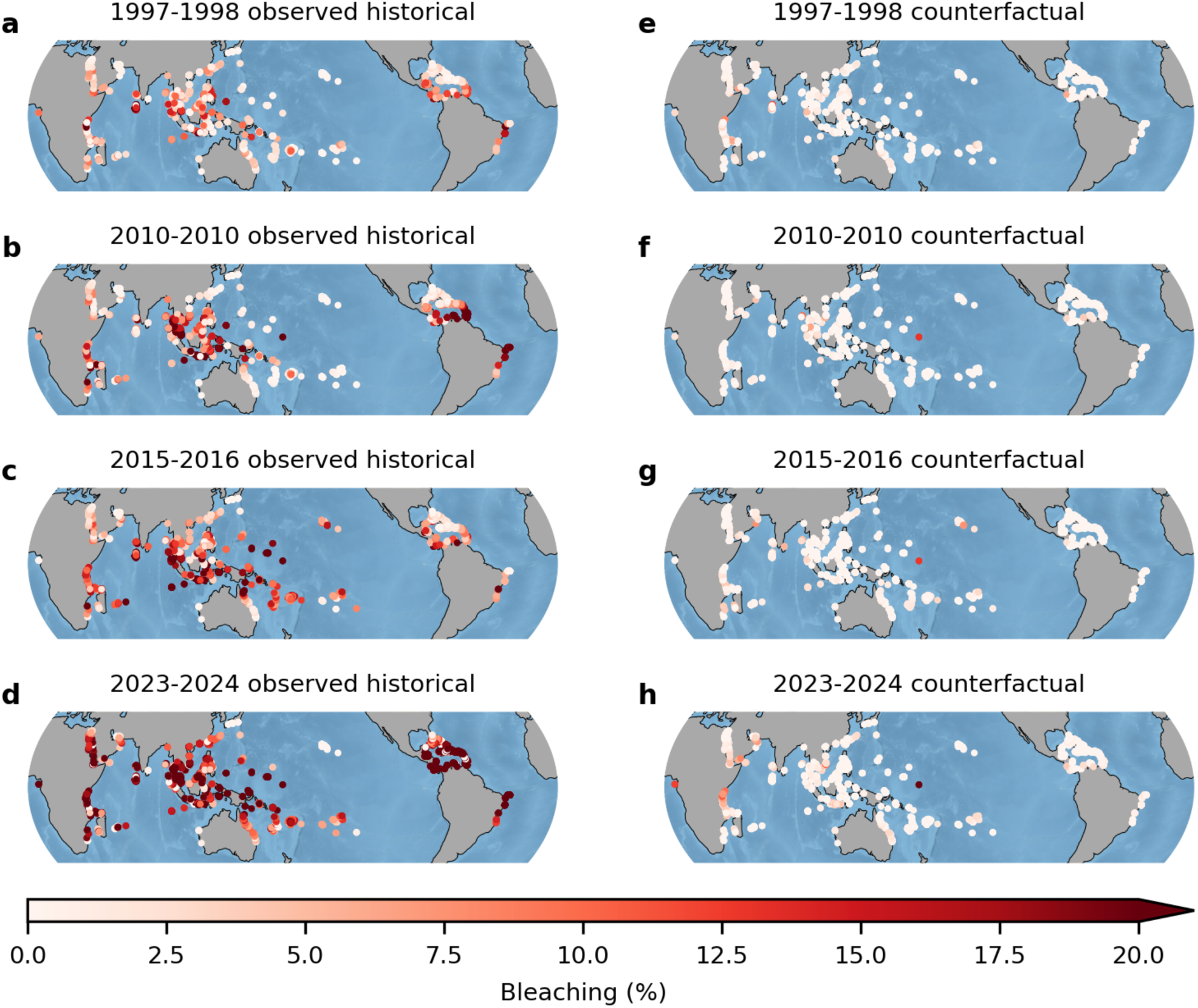
Predicted heat-driven bleaching for major El Nino events in the observed and counterfactual climates. Bleaching extent during major El Niño events as factually observed (a-d) and under a counterfactual with all anthropogenic emissions removed (e-h).

**Extended Data Figure 6.**
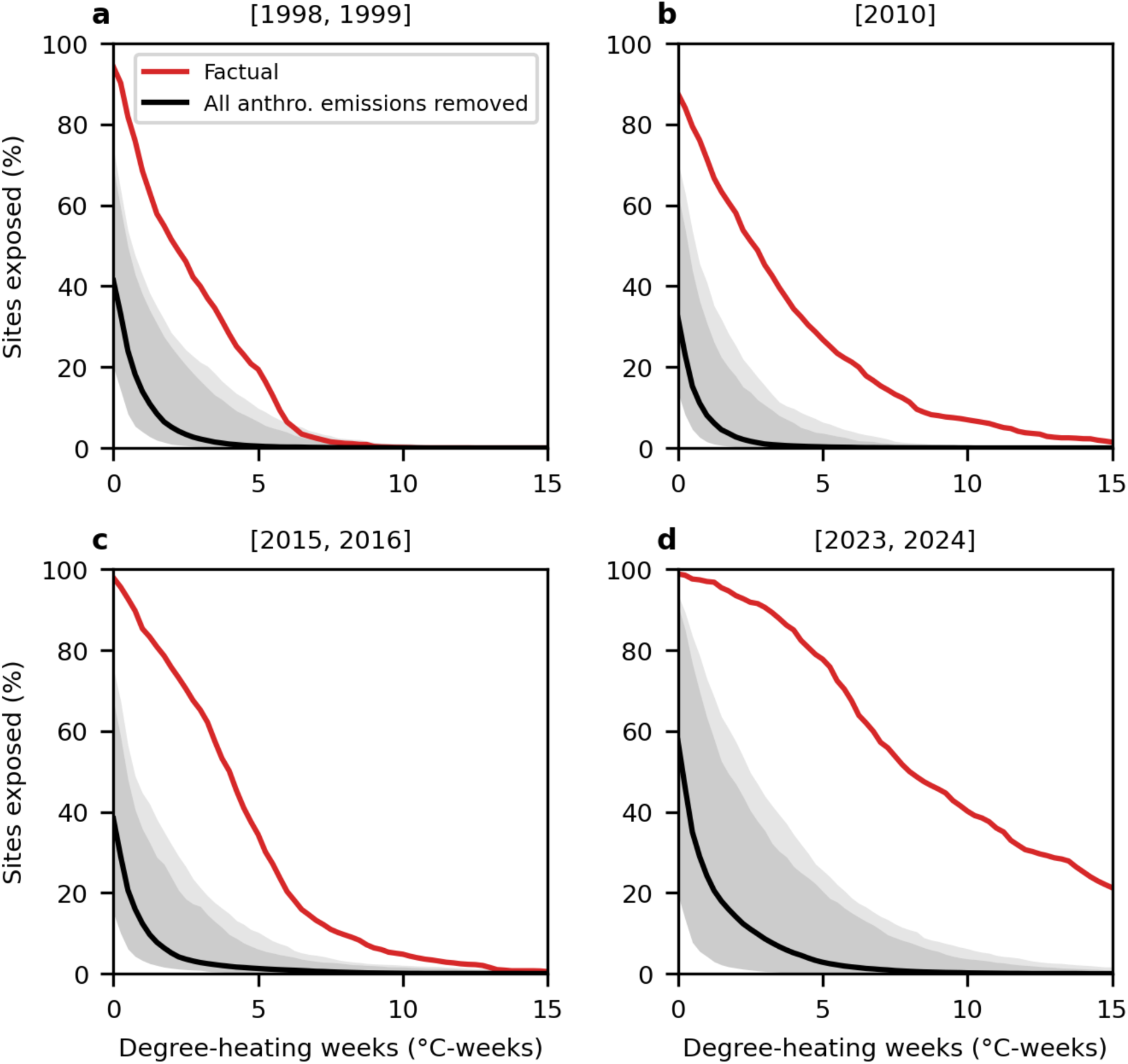
The percentage of coral reef sites exposed to increasing levels of degree-heating weeks across major El Nino events in the observed and counterfactual climates. Red curves show the factually observed number of sites exposed to a given level of degree-heating weeks, and black curves show those in counterfactual climates with all anthropogenic emissions removed. Solid curves show the median across uncertainty samples, with darker shaded areas showing the 95% (extremely likely) range, and the lighter black shading showing the upper 99th percentile of the counterfactual, which is the IPCC-based ‘virtually certain’ upper bound.

**Extended Data Figure 7.**
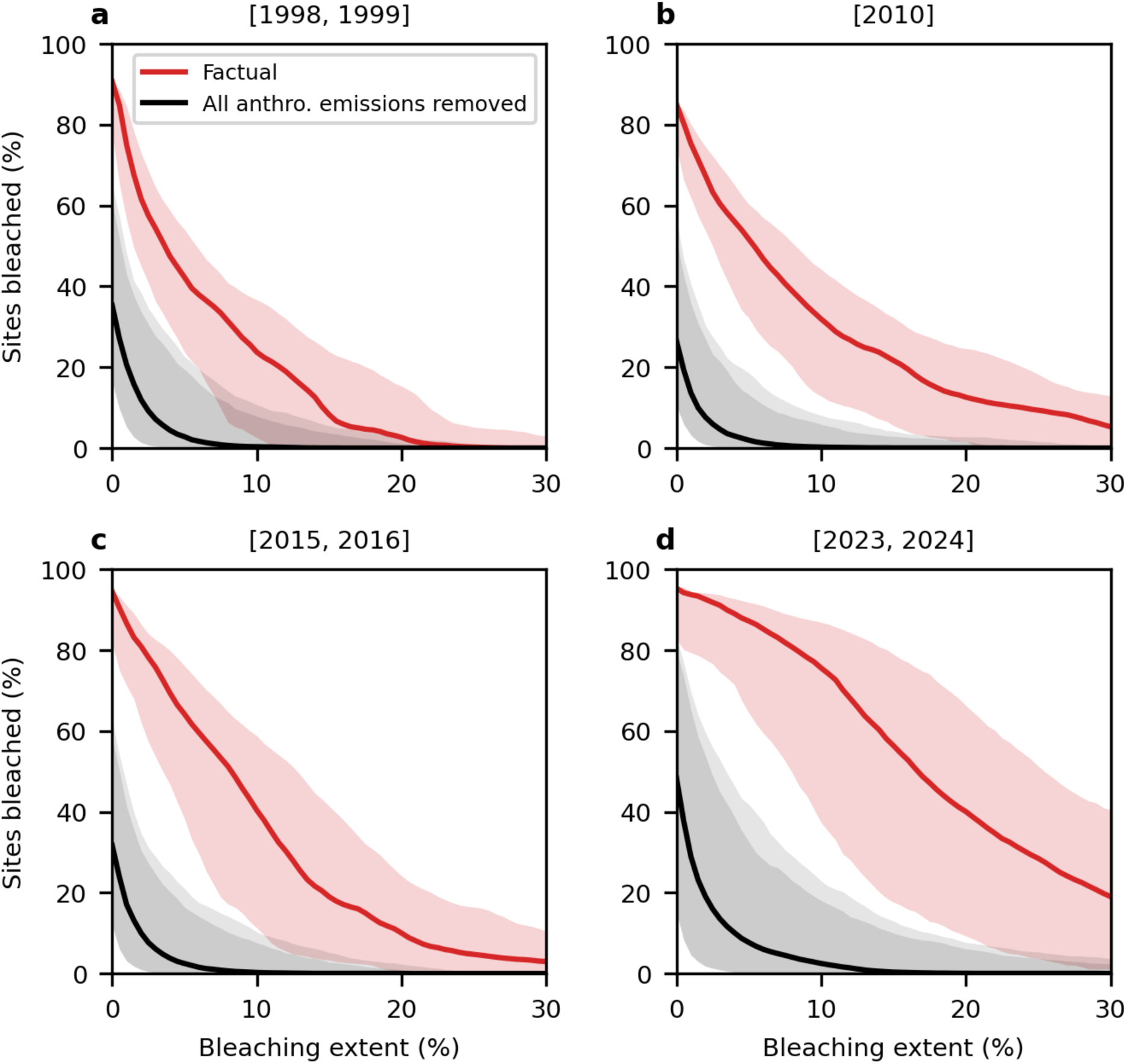
The percentage of sites exposed to increasing levels of heat-driven bleaching across major El Nino events in the observed and counterfactual climates. Red curves show the percentage of sites predicted to experience a given level of heat-driven bleaching under the observed climate and black curves show those in counterfactual climates with all anthropogenic climate forcing removed. Solid curves show the median across uncertainty samples, with darker shaded areas showing the 95% (extremely likely) range, and the lighter black shading showing the upper 99th percentile of the counterfactual, which is the IPCC-based ‘virtually certain’ upper bound.

**Extended Data Figure 8.**
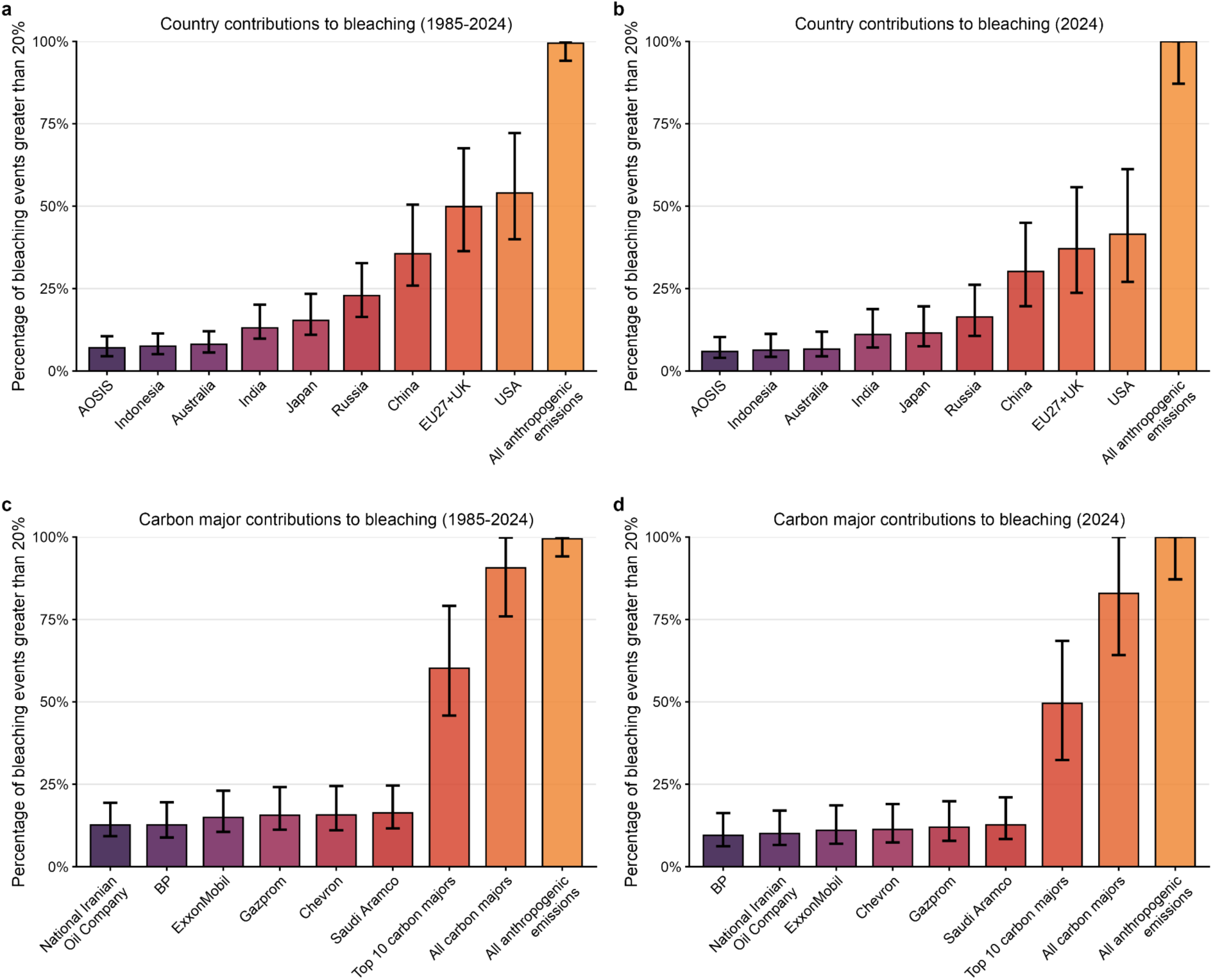
Contributions to coral reef bleaching from marine heat by states and carbon majors. As Figure 3 of the main manuscript but for bleaching of more than 20% rather than bleaching of more than 10%. **a,** The percentage of coral reef bleaching events of over 20% from 1985–2024 which would not have occurred without the cumulative, territorial emissions of individual countries or country blocs. **b,** The percentage of coral reef bleaching events of over 20% during the 2023–2024 El Niño which would not have occurred without the cumulative, territorial emissions of individual countries or country blocs. **c,d**, as for panels a and b, but showing contributions from the emissions of major emitting companies (‘carbon majors’). Bar plots are coloured based on the size of the contribution to coral bleaching and error bars show 95% confidence intervals. “All carbon majors” refers to the emissions of 172 investor or state-owned fossil fuel companies in the Carbon Majors database over the period 1850–2024. AOSIS is the 39 countries participating in the Alliance of Small Island States and EU27+UK refers to the 27 countries of the European Union and the United Kingdom. See Supplementary Tables 5–8 for values for individual states and companies.

**Extended Data Figure 9.**
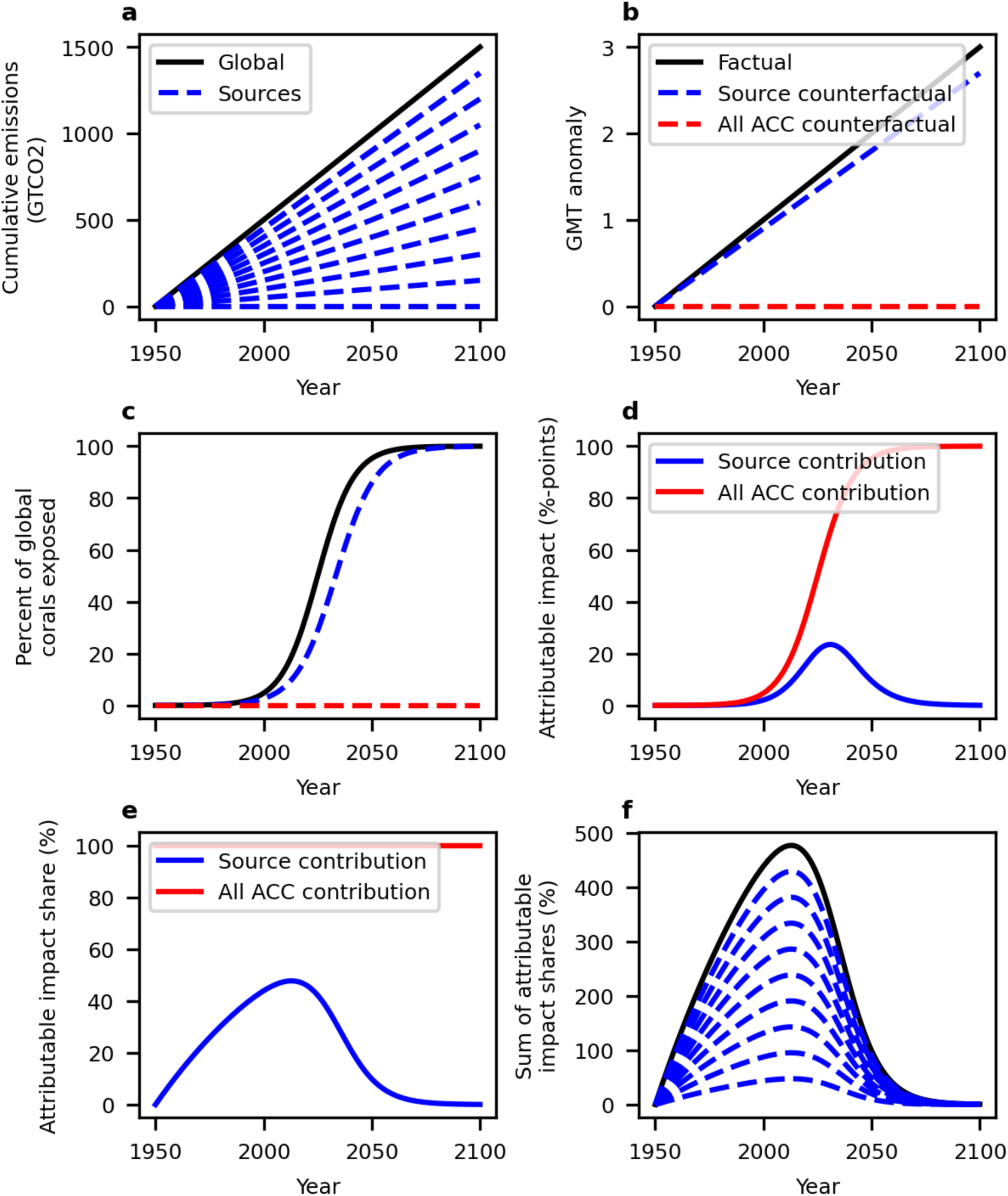
A conceptual model of source attribution with non-linear climate impacts. **a,** Cumulative emissions globally (black) and of individual emission sources (blue) - each comprising one tenth of global emissions. **b,** The global mean temperature response to cumulative emissions (assuming a climate sensitivity of 2K per 1,000GTCO2, black) and under counterfactual scenarios with the emissions of an individual source removed (blue) as well as all anthropogenic emissions removed (“ACC counterfactual”, red). **c,** The percentage of global coral reefs exposed to dangerous levels of heat under the factual and counterfactual scenarios, based on a damage curve corresponding to the saturation effect of heat exposure of global coral reefs projected to occur at levels of global warming between 1.5–2 °C. **d,** The extent of impact attributable to each counterfactual based on a “but for” framework that takes the difference between the factual and counterfactual curves from panel c. Impacts attributable to individual emission sources show a peak and decline behaviour (blue), even as impacts attributable to all anthropogenic emissions grow (red). **e,** The attributable impacts expressed as a % of all impacts. **f,** The sum of attributable impacts (black) across the ten different emission sources (blue), expressed as a share of the overall impacts, can exceed 100%.

**Extended Data Figure 10.**
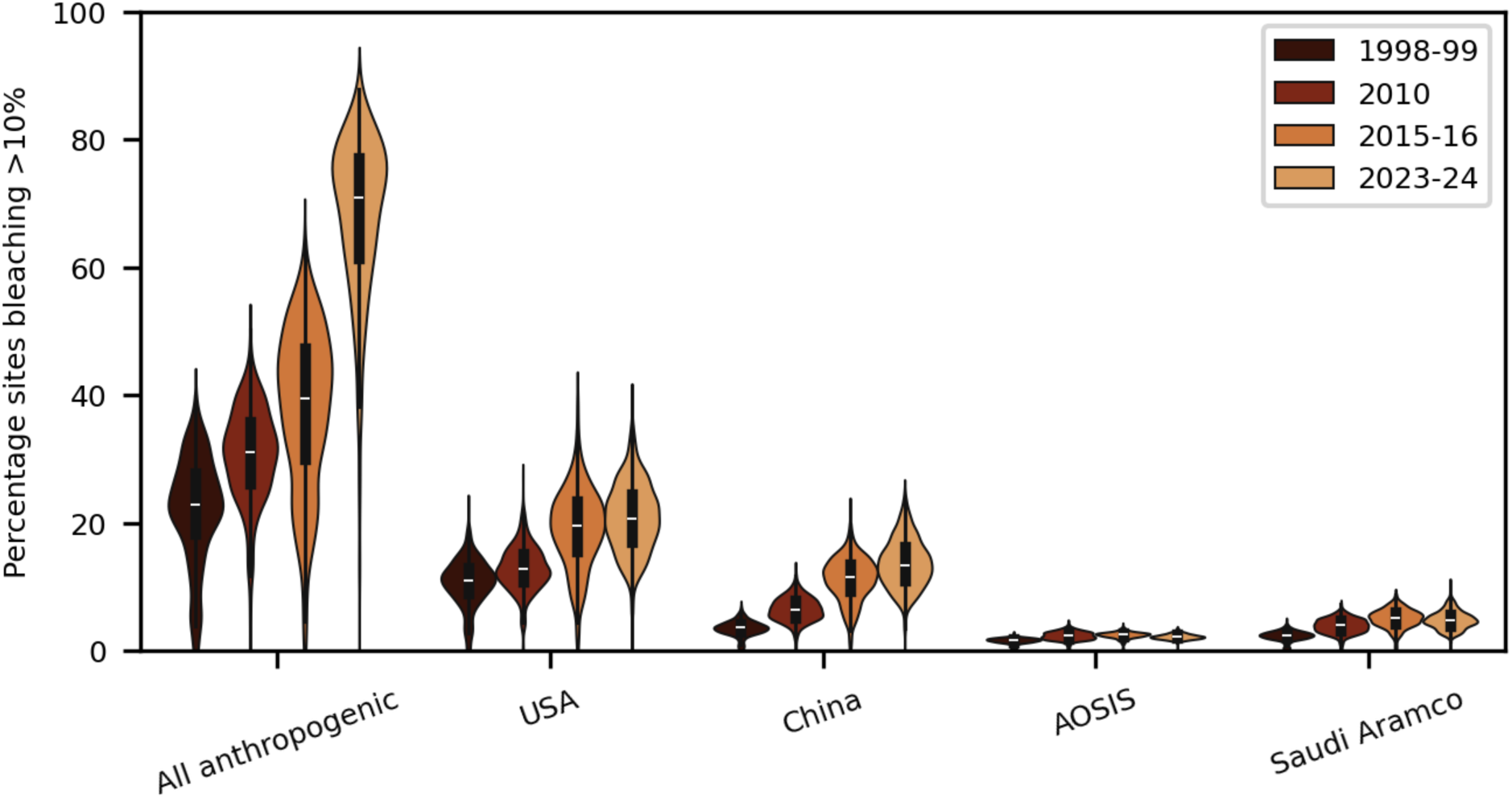
Contributions to coral reef bleaching across recent El Niño events. The percentage of sites bleaching at 10% in the observed but not in the counterfactual climate with different emissions removed are shown across major El Niño events (1998–99, 2010, 2015–16 and 2023–24). Despite large increases in the contribution of all anthropogenic emissions to bleaching between the 2015-16 and the 2023-24 event, contributions from individual sources remained constant or even declined.

## Supplementary Information

**Supplementary Figure 1.**
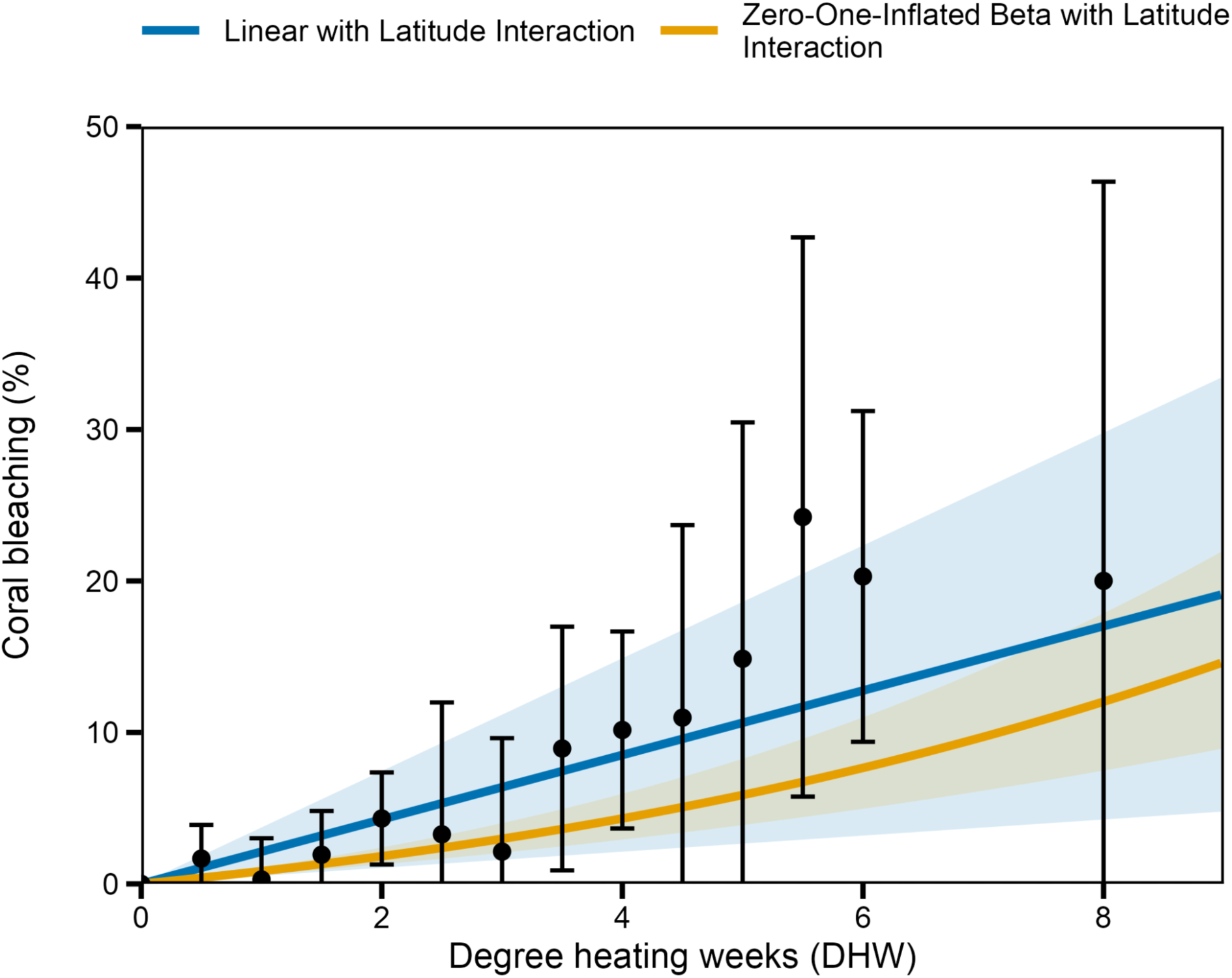
Comparison of impact model specifications. Predicted percent coral bleaching as a function of degree heating weeks (DHW) for a linear fixed-effects model (blue) and a zero-one-inflated beta regression model (orange), with 95% confidence intervals shown as shaded bands. Black points show observed mean bleaching within DHW bins of width 2°C-weeks (error bars: 95% CI across sites), providing a non-parametric reference for assessing the linearity of the DHW-bleaching relationship. Both models capture the observed DHW-bleaching relationship well across the range of the data, supporting the adequacy of the linear functional form as the preferred specification.

**Supplementary Figure 2.**
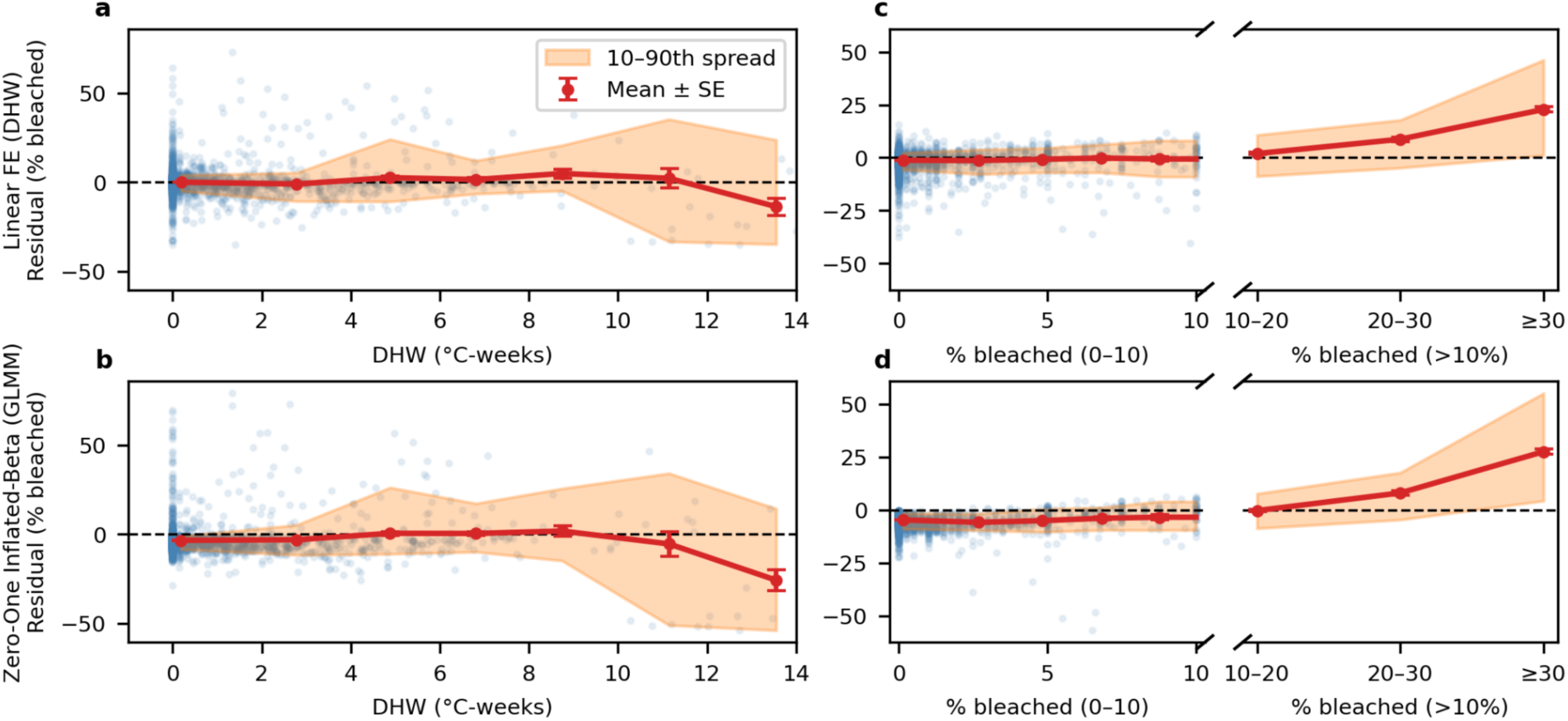
Residuals of candidate impact models as a function of DHW and bleaching. (a) residuals of the linear and (b) Zero-One-Inflated Beta GLMM model as a function of DHW. Individual residuals are shown in blue while the mean and its standard error are shown in red and the 10-90th percentile spread across different parts of the distribution of DHW (0-2, 2-4, 4-6, 6-8, 8-10, 10-12, 12-14). (c) residuals of the linear fixed-effects model and (d) the Zero-One-Inflated Beta GLMM model as a function of percentage of corals bleached across parts of its distribution (0-2, 2-4, 4-6, 6-8, 8-10, 10-20, 20-30, 30-100).

**Supplementary Figure 3.**
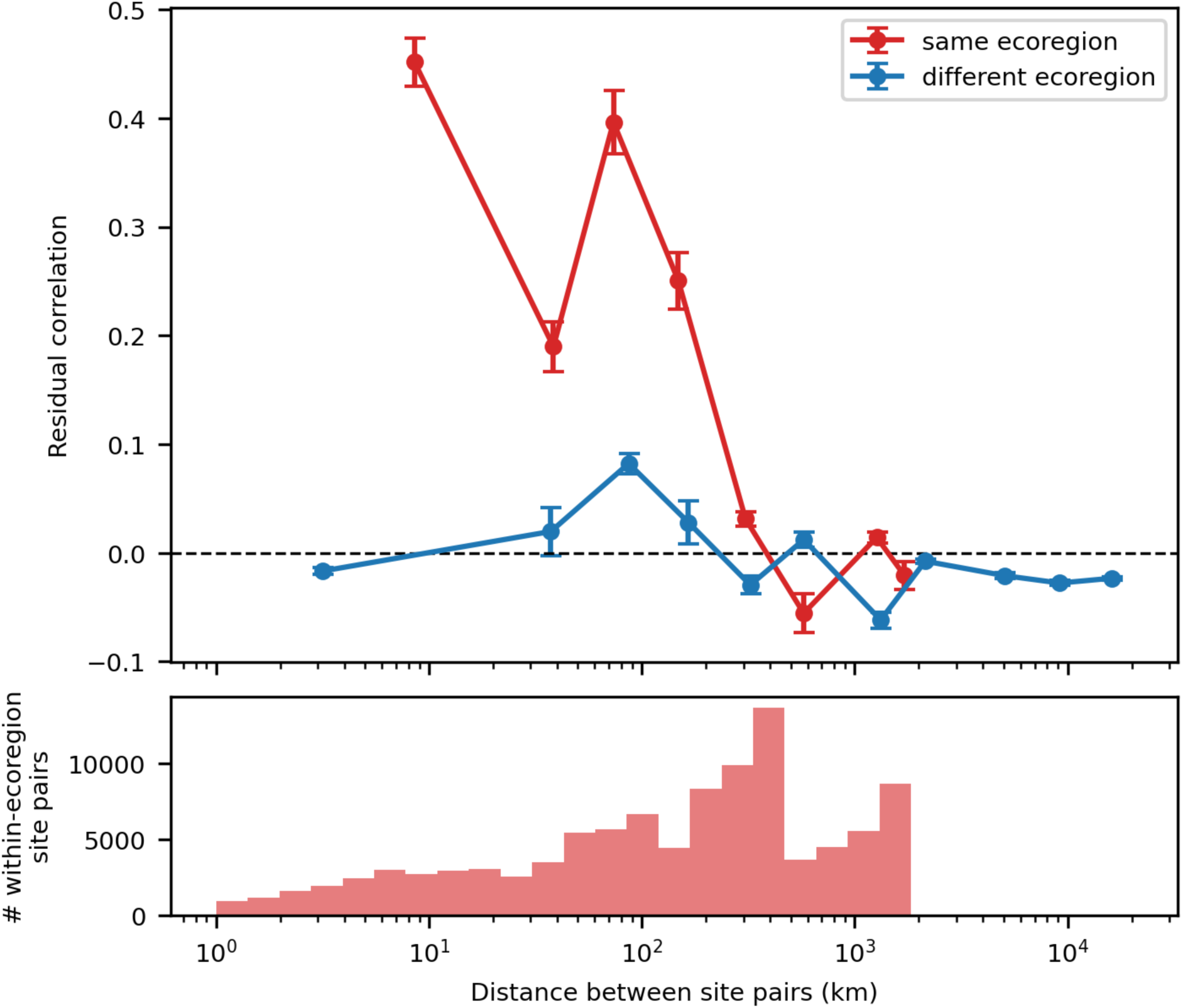
Spatial autocorrelation of linear impact model residuals. Pairwise Pearson correlation of the residuals of the Fixed-Effects linear impact model with an absolute latitude interaction (see Equation 2 in Methods) as a function of distance between pairs of coral reef sites, for sites within the same ecoregion and different ecoregions.

**Supplementary Figure 4.**
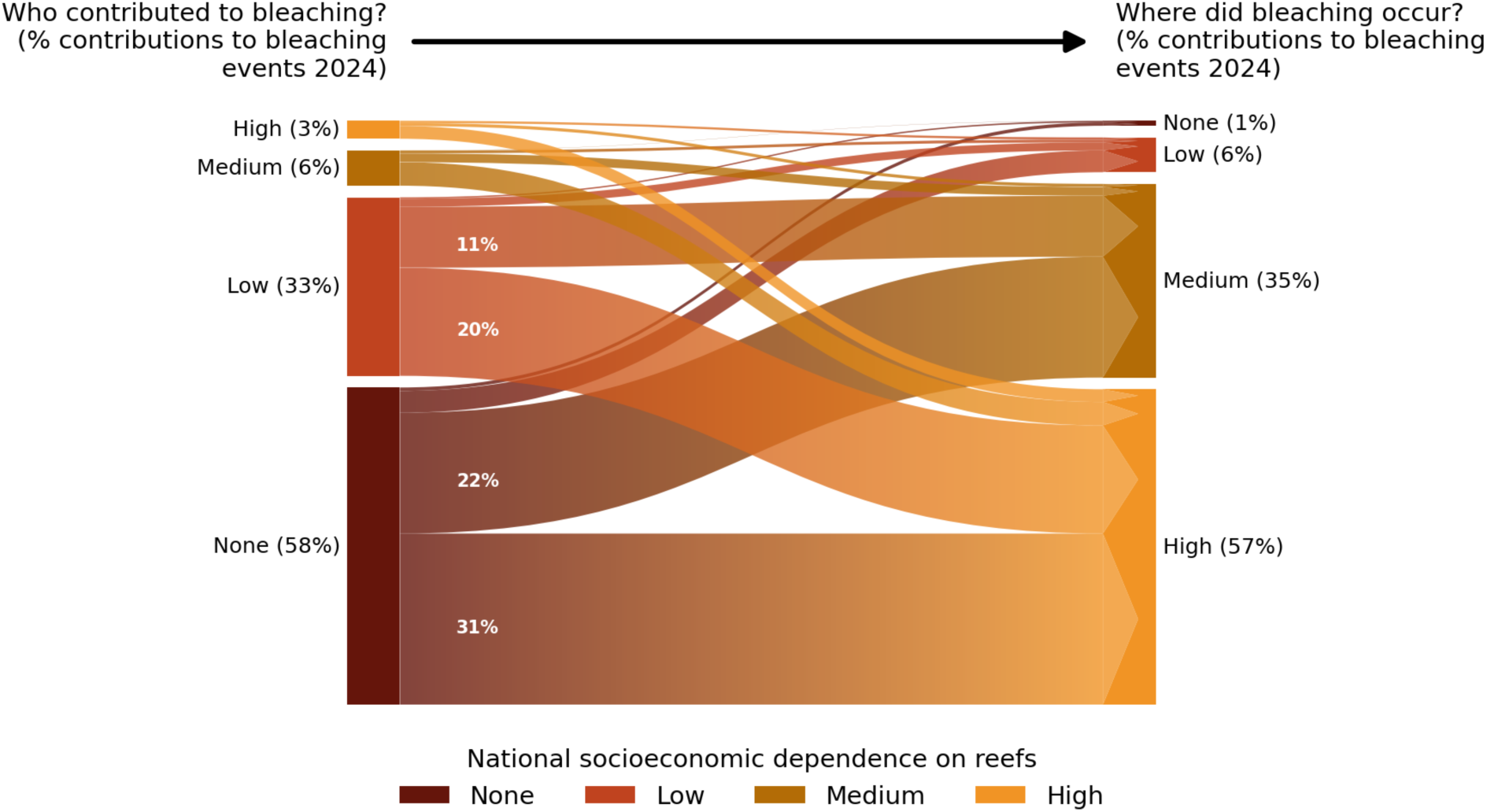
Inequalities in mass bleaching contributions and exposure during the 2024 El Niño. Same as Figure 4 of the main manuscript but for bleaching events during 2024.

**Supplementary Figure 5.**
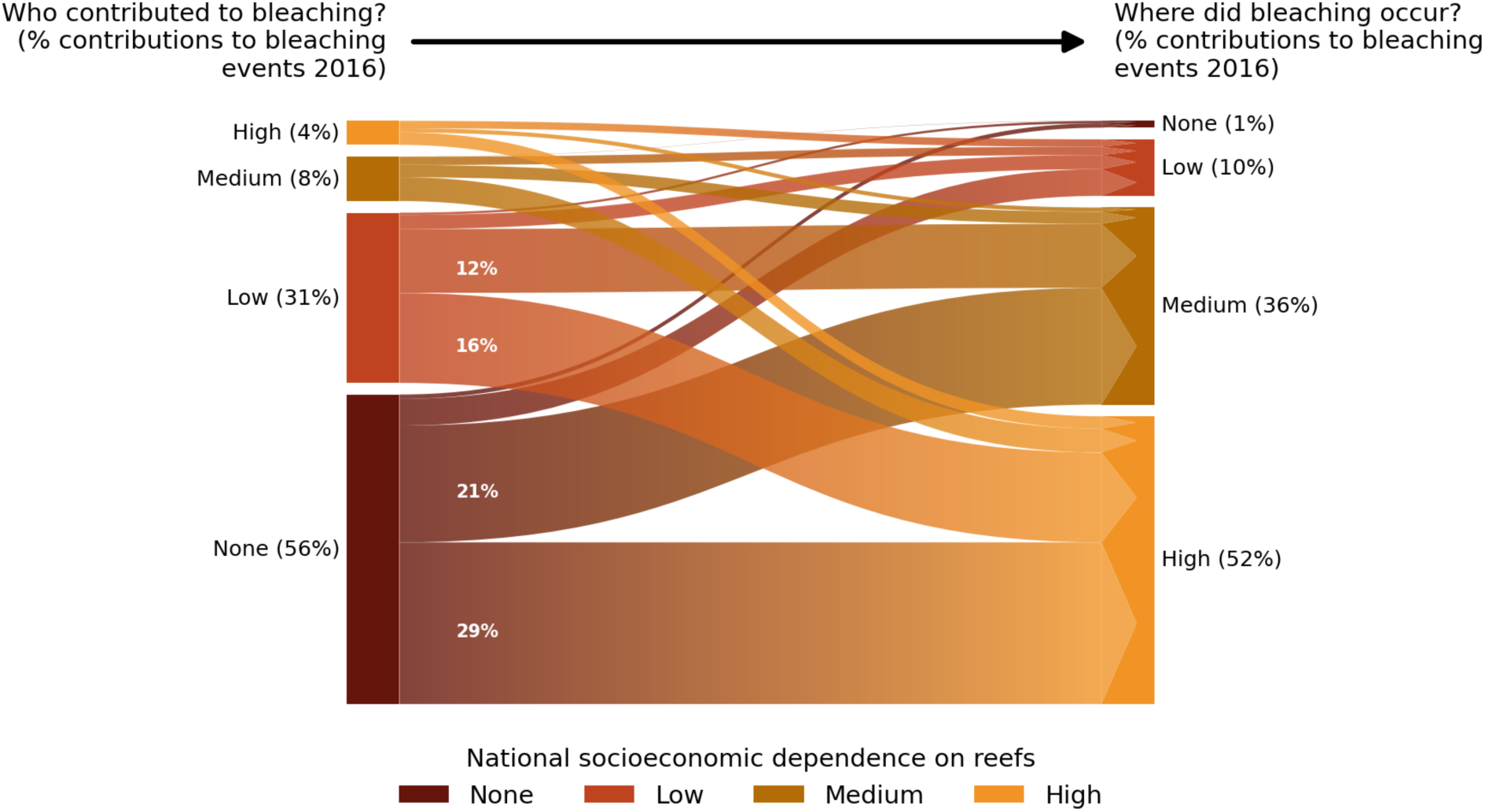
Inequalities in mass bleaching contributions and exposure during the 2016 El Niño. Same as Figure 4 of the main manuscript but for bleaching events during 2016.

**Supplementary Figure 6.**
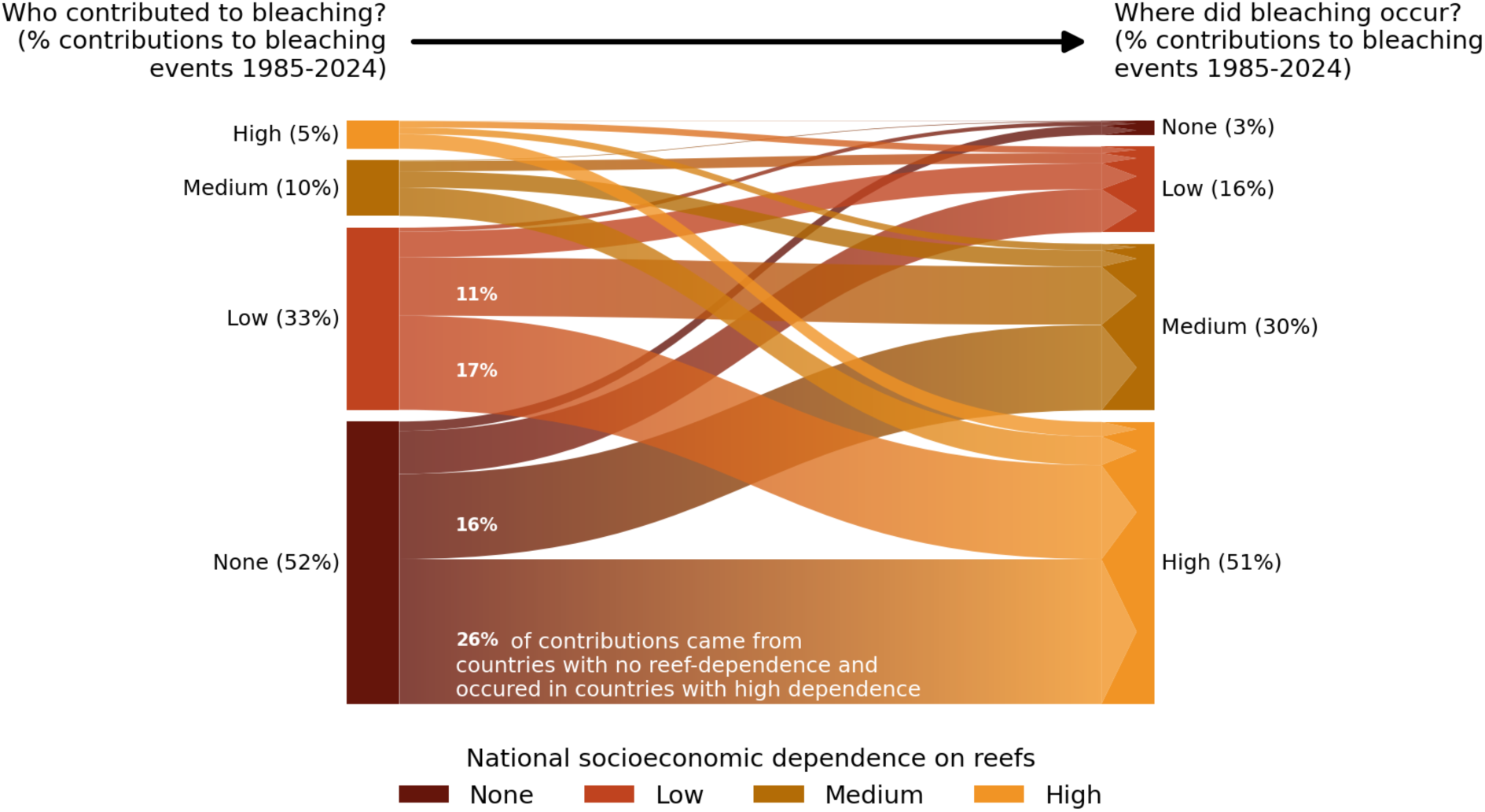
Inequalities in bleaching contributions and exposure. Same as Figure 4 of the main manuscript but for severe bleaching (>20% bleaching) rather than mass bleaching.

**Supplementary Figure 7.**
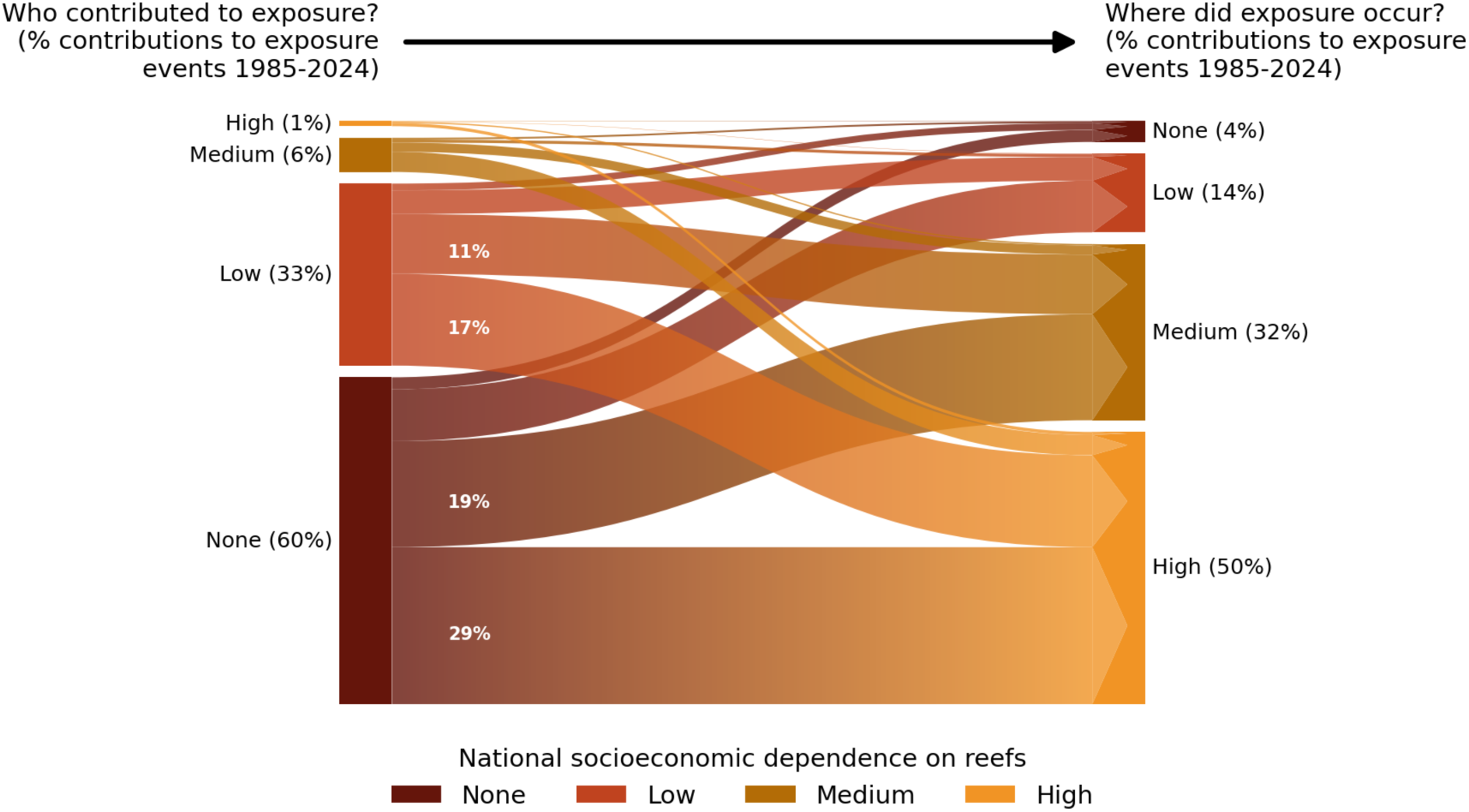
Inequalities in contributions and exposure to DHW. Same as Figure 4 of the main manuscript but for DHW exposure of 4C-weeks rather than severe bleaching.

**Supplementary Figure 8.**
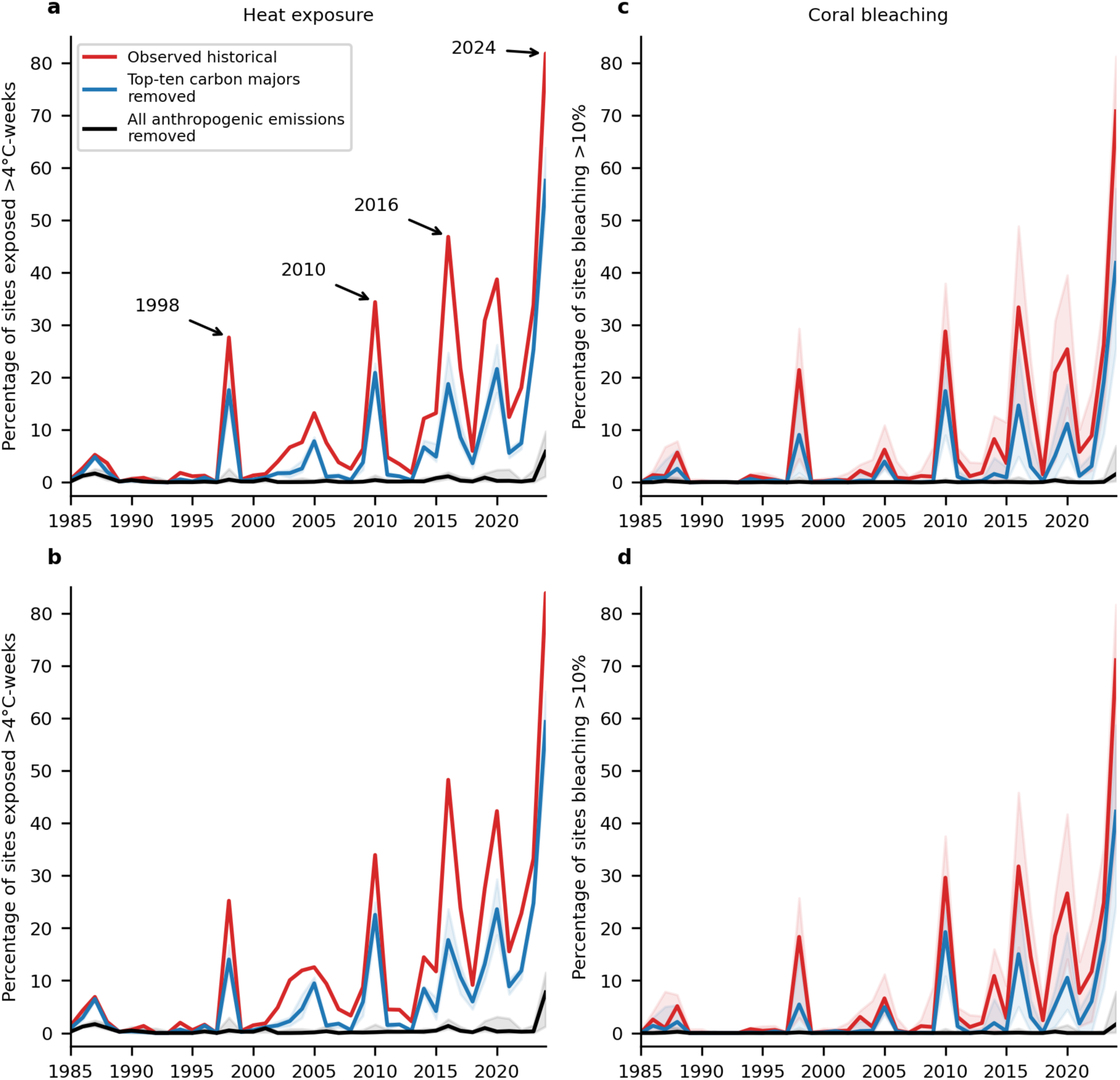
Global coral reef exposure and bleaching attribution with a reduced set of coral sites. (a) Observed and counterfactual historical heat exposure and (b) coral bleaching across all coral sites as in Figure 2a and d, (b, d) as well as for the smaller set of sites with repeated bleaching observations used to estimate the impact model shown in Figure 1b.

**Supplementary Figure 9.**
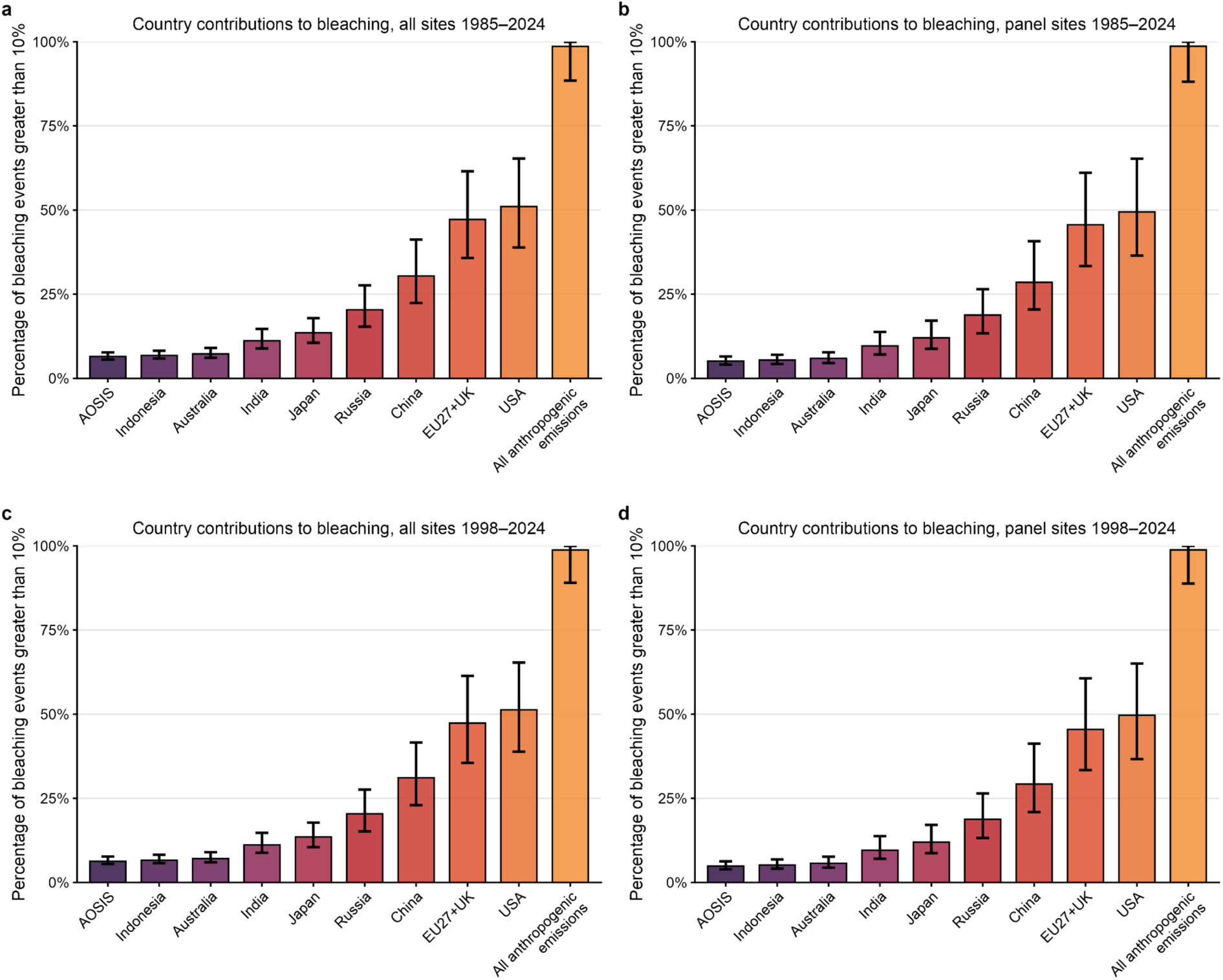
Robustness of estimated coral bleaching from marine heat by countries and companies to changes in numbers of sites and selection of time periods. Each panel shows the percentage of site-level bleaching events (>10% bleaching) attributable to individual countries’ historical emissions, and to all anthropogenic emissions combined. Panels compare results for the full set of sites in the coral reef survey data against the panel regression subset of sites (sites with repeated observations) used in the DHW–bleaching impact model, as well as the 1985–2024 period against the 1998-2024 period: (**a**) all sites 1985–2024; (**b**) panel regression sites 1985–2024; (**c**) all sites 1998–2024; (**d**) panel regression sites 1998–2024. Bars show median attribution across bootstrap samples; error bars show 95% confidence intervals. Country ordering and attribution estimates are highly consistent across all four specifications, indicating the results are robust to the choice of site sample and time period.

**Supplementary Figure 10.**
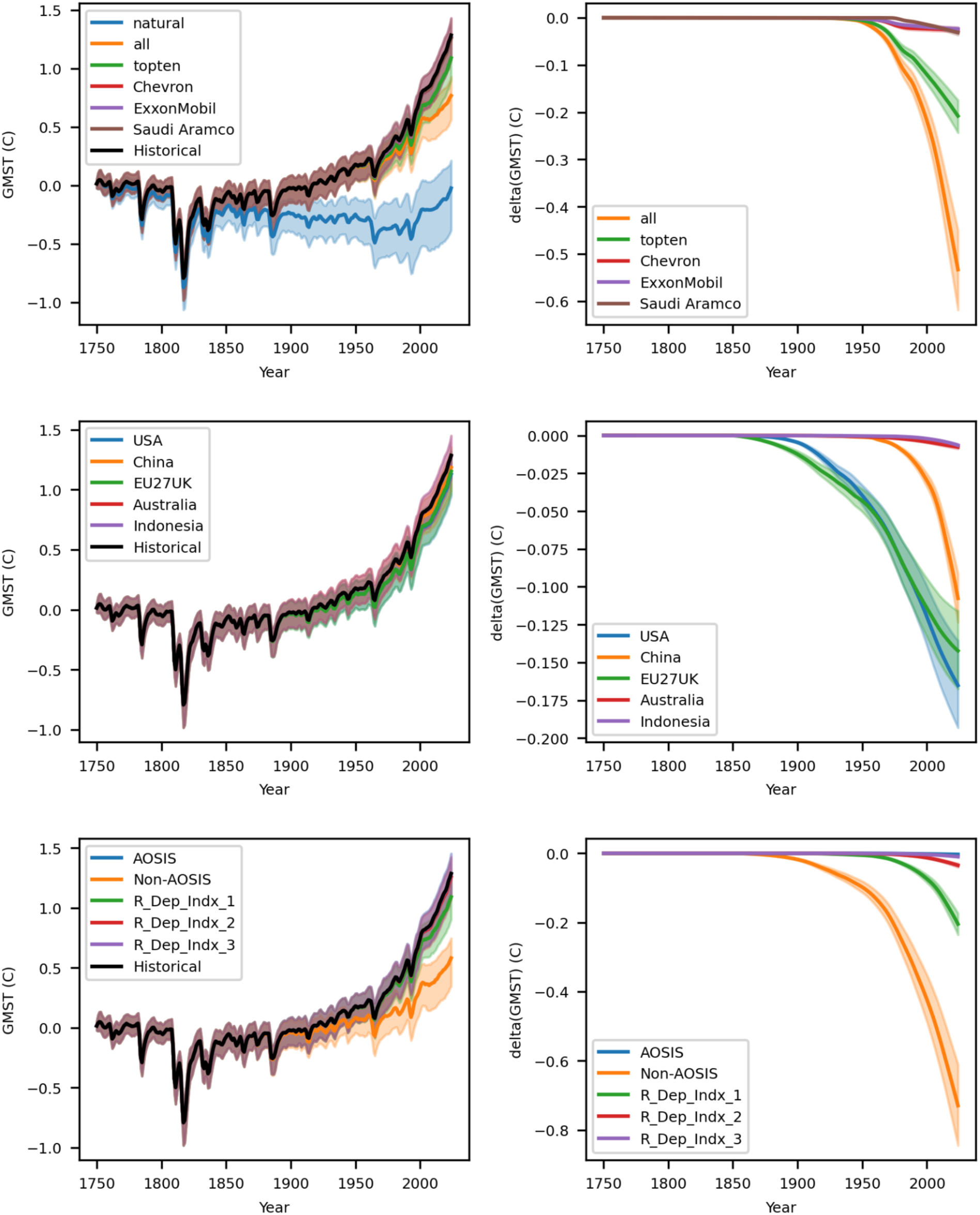
Global temperature counterfactuals simulated by FaIR. Global mean surface temperature (GMST) simulated by FaIR (left-hand column) under scenarios removing company (upper), country (middle) and groups of countries by reef dependence index or membership of AOSIS (lower) are shown. “Historical” indicates the historically observed emission scenario with no emissions removed, and “natural” the counterfactual with all anthropogenic emissions removed. “All” indicates the removal of all carbon majors, “topten”, “topfive” and “All_USA” the removal of the top ten top five and all USA based carbon majors. The difference in GMST between these counterfactual scenarios and the factual scenario with no emissions removed are shown (right hand column).

**Supplementary Figure 11.**
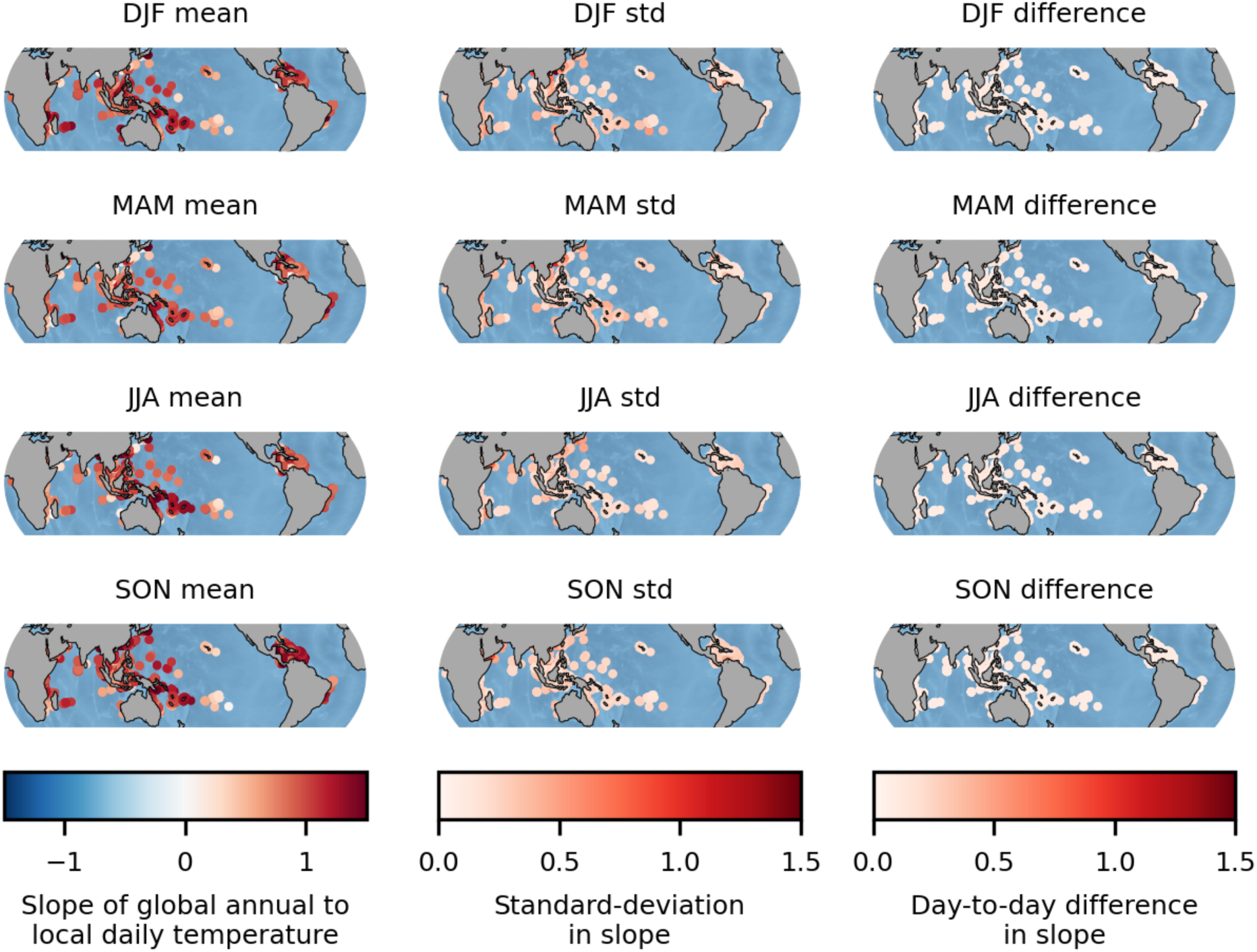
Seasonal averages, standard deviations, and day-to-day differences of the pattern scaling factor between global temperature and local daily temperature. The pattern correlations between global mean surface temperature and local daily sea surface temperature are averaged across days within a season (left), and their standard deviation is shown (central column), along with the average absolute difference of scaling factors between adjacent calendar days of the year (right hand column).

**Supplementary Figure 12.**
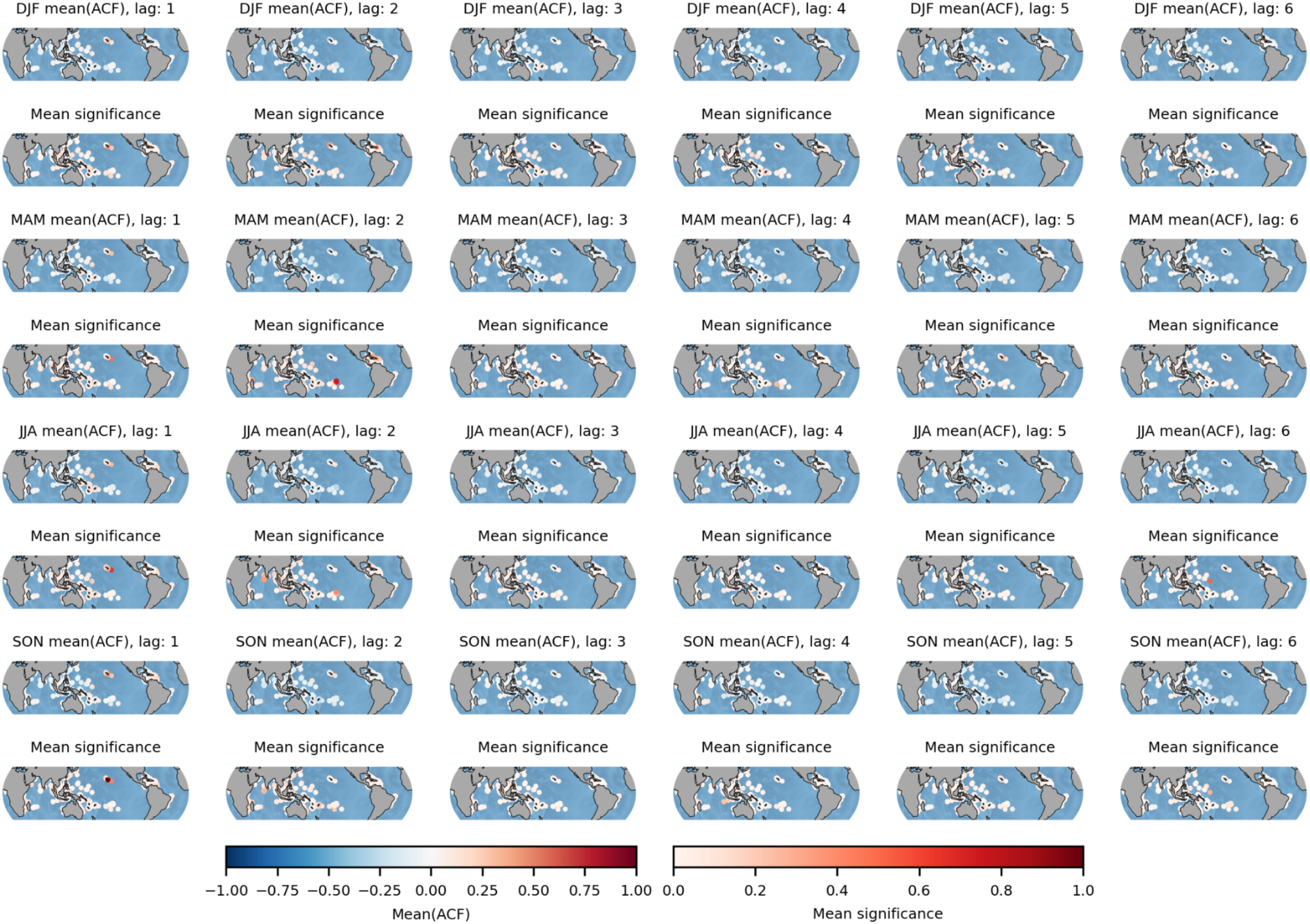
Seasonal averages and significance of the autocorrelation function (ACF) of the residual between global and local daily temperatures. The autocorrelation of the residuals between the global annual mean surface temperature and local daily sea surface temperatures are calculated for each day of the year and grid-cell site and their significance is tested with respect to a 5% significance level using Bartlett’s formula. The mean average ACF and mean significance of these ACFs across calendar days within individual seasons (ordered top to bottom) are shown at lags of 1 to 6 years (left to right).

**Supplementary Figure 13.**
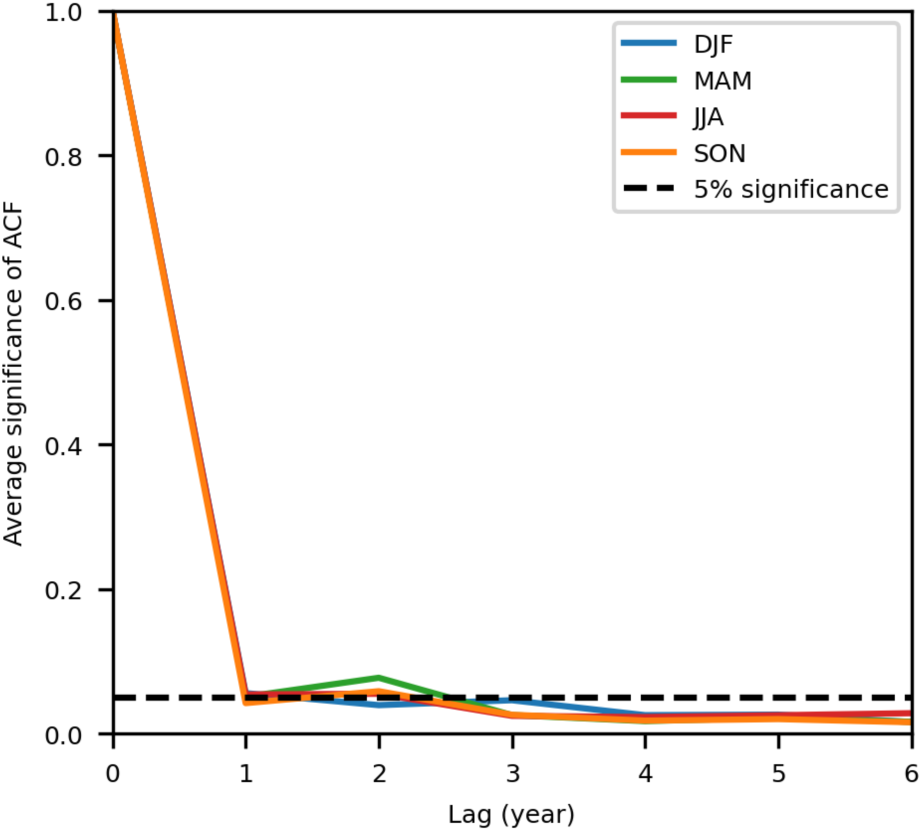
Average significance of autocorrelation function (ACF) of pattern scaling residuals. The autocorrelation of the residuals between the global annual mean surface temperature and local daily sea surface temperatures are calculated for each day of the year and grid-cell site, their significance is tested with respect to a 5% significance level using Bartlett’s formula, and the average number of significant autocorrelations is taken across sites and days of the year within a given season. The expected number of significant correlations is shown as the horizontal dashed black line. The strong decay of significance in the autocorrelation functions with the first year’s lag indicates a very weak autocorrelation structure of residuals across years.

**Supplementary Figure 14.**
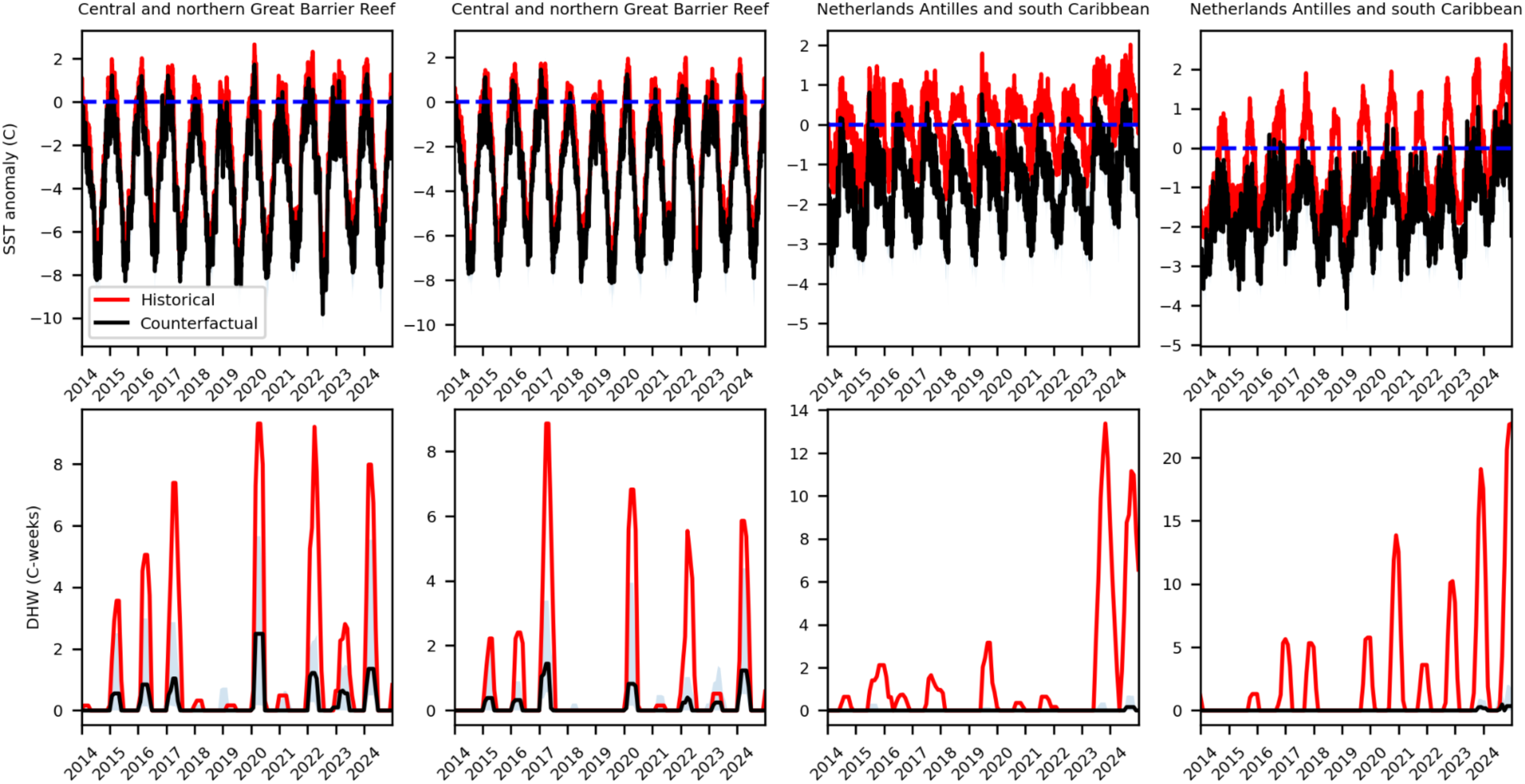
Observed and counterfactual historical sea surface temperature anomalies and degree-heating weeks from exemplary sites. The upper row shows Sea Surface Temperature (SST) anomalies relative to the threshold used to define degree-heating week exposure (dashed blue line) for two sites in two ecoregions under the observed historical (red) and counterfactual historical scenario with all anthropogenic emissions removed (black). The lower row shows Degree-Heating-Weeks calculated from these SST anomalies.

**Table S1.** Testing interaction terms for heterogeneities in bleaching sensitivity to DHWs. All columns show results for panel regression models with percentage bleached as the dependent variable. All models include site, year and seasonality(ecoregion x month) fixed effects, with standard errors accounting for spatial autocorrelation using the Conley (200 km) method. Different columns indicate the inclusion of different variables as an interaction term with DHW to test heterogeneity in the coral bleaching response to DHW across sites. DHW and SST anomaly variability indicate the historical (1985-2005) standard deviation of DHW or SST anomalies. SST anomaly warming indicates the rate of warming of SST anomalies based on the difference in the average SST values prior to and after 2005.

|  | *Dependent variable: Percent Bleached (%)* |  |  |  |  |  |  |  |
| --- | --- | --- | --- | --- | --- | --- | --- | --- |
|  | Latitude Interaction<br>(1) | Turbidity Interaction<br>(2) | Depth Interaction<br>(3) | DHW Anomaly<br>Variability Interaction<br>(4) | SST Anomaly<br>Variability Interaction<br>(5) | SST Anomaly<br>Warming Interaction<br>(6) | Latitude and SST<br>Anomaly Variability Interaction<br>(7) | Marine Protected Area<br>Interaction<br>(8) |
| DHW | 3.82**<br>(1.09) | 2.50**<br>(0.81) | 2.64*<br>(1.06) | 2.50<br>(1.51) | 3.25**<br>(1.03) | 2.69<br>(1.80) | 3.82**<br>(1.15) | 1.28**<br>(0.35) |
| DHW ×<br>\|Latitude\| | −0.13**<br>(0.05) |  |  |  |  |  | −0.13<br>(0.07) |  |
| DHW ×<br>Turbidity |  | −6.63<br>(3.85) |  |  |  |  |  |  |
| DHW ×<br>Depth |  |  | −0.11<br>(0.09) |  |  |  |  |  |
| DHW ×<br>DHW Anomaly |  |  |  | −0.90<br>(1.33) |  |  |  |  |
| DHW ×<br>SST Anomaly Variability |  |  |  |  | −0.91*<br>(0.39) |  | 0.00<br>(0.69) |  |
| DHW ×<br>SST Anomaly Warming |  |  |  |  |  | −2.05<br>(3.81) |  |  |
| DHW ×<br>Marine Protected Area |  |  |  |  |  |  |  | 0.76<br>(0.68) |
| Observations | 6897 | 6491 | 6491 | 6897 | 6897 | 6897 | 6897 | 6897 |
| $R^2$ | 0.582 | 0.468 | 0.468 | 0.574 | 0.578 | 0.574 | 0.582 | 0.575 |
| $R^2$ Within | 0.071 | 0.079 | 0.077 | 0.054 | 0.062 | 0.053 | 0.071 | 0.054 |
| BIC | 65 101.5 | 59 080.6 | 59 091.7 | 65 227.7 | 65 169.3 | 65 233.8 | 65 110.3 | 65 222.0 |

\* p < 0.05, \*\* p < 0.01

**Table S2.** Effects of different order polynomials on DHW for bleaching. All columns show results for panel regression models with percentage bleached as the dependent variable. All models include site, year, and seasonality(ecoregion x month) fixed effects, with standard errors accounting for spatial autocorrelation using the Conley (200 km) method. Sensitivity testing of the linear latitude model with 2nd and 3rd degree polynomials are presented in this table. The 3rd degree polynomial (cubic) model does not display any significant terms, whereas, the 2nd degree polynomial (quadratic) does but not for the latitude interaction term. Thus, the 1st degree polynomial remains the most suitable model for the data.

| *Dependent variable: Percent Bleached (%)* |  |  |  |  |  |  |
| --- | --- | --- | --- | --- | --- | --- |
|  | Linear with Latitude Interaction<br>(1) | Linear<br>(2) | Quadratic with Latitude Interaction<br>(3) | Quadratic<br>(4) | Cubic with Latitude Interaction<br>(5) | Cubic<br>(6) |
| DHW | 3.82**<br>(1.09) | 1.76**<br>(0.57) | 3.63<br>(1.85) | 2.59*<br>(1.15) | 1.96<br>(2.38) | 1.07<br>(1.29) |
| DHW × \|Latitude\| | −0.13**<br>(0.05) |  | −0.05<br>(0.12) |  | −0.12<br>(0.12) |  |
| DHW<sup>2</sup> |  |  | 0.05<br>(0.23) | −0.10<br>(0.14) | 0.49<br>(0.75) | 0.32<br>(0.36) |
| DHW<sup>2</sup> × \|Latitude\| |  |  | −0.01<br>(0.02) |  | 0.01<br>(0.04) |  |
| DHW<sup>3</sup> |  |  |  |  | −0.02<br>(0.05) | −0.02<br>(0.02) |
| DHW<sup>3</sup> × \|Latitude\| |  |  |  |  | 0.00<br>(0.00) |  |
| Observations | 6897 | 6897 | 6897 | 6897 | 6897 | 6897 |
| R<sup>2</sup> | 0.582 | 0.573 | 0.584 | 0.575 | 0.590 | 0.577 |
| R<sup>2</sup> Within | 0.071 | 0.052 | 0.076 | 0.055 | 0.088 | 0.060 |
| BIC | 65 101.5 | 65 233.8 | 65 081.9 | 65 215.3 | 65 009.3 | 65 192.4 |

\* p < 0.05, \*\* p < 0.01

**Table S3.** Testing the robustness of the impact model to alternative DHW definitions. All columns show results for panel regression models with percentage bleached as the dependent variable. All models include site, year and seasonality(ecoregion x month) fixed effects, with standard errors accounting for spatial autocorrelation using the Conley (200 km) method. DHW values are accumulated over 12 weeks and based on exceedances of 1 °C of the maximum monthly climatology. Adjusted DHW values are accumulated over 11 rather than 12 weeks and based on exceedances of 0.4 °C rather than 1 °C of the maximum monthly climatology, as suggested by Whitaker et. al^1^.

|  | *Dependent variable: Percent Bleached (%)* |  |  |  |
| --- | --- | --- | --- | --- |
|  | DHW with<br>Latitude Interaction<br>(1) | DHW<br>(2) | Adjusted DHW<br>(3) | Adjusted DHW with<br>Latitude Interaction<br>(4) |
| DHW | 3.82**<br>(1.09) | 1.76**<br>(0.57) |  |  |
| DHW × \|Latitude\| | −0.13**<br>(0.05) |  |  |  |
| Adjusted DHW |  |  | 1.47**<br>(0.36) | 2.67**<br>(0.67) |
| Adjusted DHW × \|Latitude\| |  |  |  | −0.08*<br>(0.03) |
| Observations | 6897 | 6897 | 6897 | 6897 |
| $R^2$ | 0.582 | 0.573 | 0.578 | 0.584 |
| $R^2$ Within | 0.071 | 0.052 | 0.062 | 0.075 |
| BIC | 65 101.5 | 65 233.8 | 65 159.3 | 65 068.3 |

\* $p < 0.05$ , \*\* $p < 0.01$

**Table S4.** Robustness tests across different error clustering schemes. All columns show results for panel regression models with percentage bleached as the dependent variable. All models include site, year and seasonality (ecoregion x month) fixed effects, with the relevant spatial clustering scheme indicated in each column header. Clustering by site accounts for pseudo-replication driven by temporal autocorrelation within a site; clustering by ecoregion additionally accounts for spatio-temporal autocorrelation between sites within the same ecoregion; and Conley (200 km) standard errors account for spatial autocorrelation between neighbouring sites as a continuous function of geographic distance.

| *Dependent variable: Percent Bleached (%)* |  |  |  |  |
| --- | --- | --- | --- | --- |
|  | Conley 200km Clustering<br>(1) | Ecoregion Clustering<br>(2) | Site Clustering<br>(3) | No Clustering<br>(4) |
| DHW | 3.82**<br>(1.09) | 3.82**<br>(0.81) | 3.82**<br>(0.46) | 3.82**<br>(0.23) |
| DHW × \|Latitude\| | -0.13**<br>(0.05) | -0.13**<br>(0.04) | -0.13**<br>(0.02) | -0.13**<br>(0.01) |
| Observations | 6897 | 6897 | 6897 | 6897 |
| $R^2$ | 0.582 | 0.582 | 0.582 | 0.582 |
| $R^2$ Within | 0.071 | 0.071 | 0.071 | 0.071 |
| BIC | 65 101.5 | 65 101.5 | 65 101.5 | 65 101.5 |

\* $p < 0.05$ , \*\* $p < 0.01$

**Table S5.** Percentage of events with coral bleaching greater than 10% which are attributed to the emissions of countries and companies from 1985–2024. Columns show the fraction of bleaching events at coral reef sites globally and within specific countries or country blocs that can be attributed to emissions from specific companies and countries (listed in the rows). Values are medians and uncertainty estimates are 95% confidence intervals. Asterisks (*) denote emitters for which the attributed fraction is positive across the entire bootstrap distribution (1st percentile > 0%), indicating a virtually certain contribution to bleaching in that region. Top 10 refers to the top 10 major carbon emitting companies as defined in the Carbon Majors Database ^2^, AOSIS refers to the intergovernmental coalition the Alliance of Small Island States, and EU27+UK refers to the 27 countries of the European Union and the United Kingdom. *N* denotes the number of coral reef sites present in the data for each country or country bloc.

| Scenario | Global (N=4471) | AOSIS (N=311) | Indonesia (N=89) | Japan (N=21) | China (N=65) | USA (N=25) | Australia (N=154) |
| --- | --- | --- | --- | --- | --- | --- | --- |
| All anthropogenic | 99%*<br>(88% - 100%) | 99%*<br>(91% - 100%) | 100%*<br>(89% - 100%) | 100%*<br>(76% - 100%) | 100%*<br>(100% - 100%) | 100%*<br>(91% - 100%) | 100%*<br>(82% - 100%) |
| All carbon majors | 84%*<br>(69% - 96%) | 79%*<br>(63% - 97%) | 87%*<br>(72% - 100%) | 100%*<br>(40% - 100%) | 100%*<br>(80% - 100%) | 68%*<br>(32% - 100%) | 93%*<br>(60% - 100%) |
| Saudi Aramco | 14%*<br>(11% - 19%) | 13%*<br>(7% - 20%) | 10%*<br>(3% - 17%) | 5%<br>(0% - 100%) | 10%<br>(1% - 88%) | 7%<br>(0% - 22%) | 12%*<br>(3% - 35%) |
| ExxonMobil | 14%*<br>(11% - 18%) | 13%*<br>(7% - 19%) | 10%*<br>(3% - 17%) | 5%<br>(0% - 100%) | 9%<br>(0% - 86%) | 7%<br>(0% - 21%) | 11%<br>(3% - 30%) |
| Chevron | 14%*<br>(11% - 19%) | 14%*<br>(7% - 20%) | 11%*<br>(3% - 18%) | 5%<br>(0% - 100%) | 10%<br>(0% - 86%) | 7%<br>(0% - 22%) | 11%<br>(4% - 32%) |
| Holcim Group | 6%*<br>(5% - 8%) | 4%*<br>(2% - 6%) | 3%*<br>(0% - 6%) | 0%<br>(0% - 50%) | 1%<br>(0% - 23%) | 0%<br>(0% - 6%) | 2%<br>(0% - 12%) |
| Top 10 | 56%*<br>(43% - 70%) | 55%*<br>(32% - 69%) | 55%*<br>(35% - 78%) | 46%*<br>(10% - 100%) | 73%*<br>(32% - 100%) | 31%*<br>(4% - 75%) | 67%*<br>(32% - 92%) |
| Gazprom | 14%*<br>(11% - 18%) | 13%*<br>(7% - 19%) | 9%*<br>(3% - 16%) | 5%<br>(0% - 100%) | 10%<br>(0% - 86%) | 7%<br>(0% - 21%) | 11%*<br>(3% - 34%) |
| National Iranian Oil Company | 11%*<br>(9% - 14%) | 10%*<br>(5% - 14%) | 7%*<br>(2% - 13%) | 3%<br>(0% - 84%) | 7%<br>(0% - 86%) | 4%<br>(0% - 17%) | 8%<br>(2% - 27%) |
| BP | 11%*<br>(9% - 15%) | 10%*<br>(6% - 15%) | 8%*<br>(2% - 14%) | 3%<br>(0% - 67%) | 8%<br>(0% - 82%) | 5%<br>(0% - 18%) | 8%<br>(2% - 25%) |
| China | 30%*<br>(22% - 41%) | 31%*<br>(18% - 42%) | 24%*<br>(14% - 49%) | 18%<br>(0% - 100%) | 37%*<br>(10% - 100%) | 18%<br>(0% - 38%) | 34%*<br>(13% - 67%) |
| USA | 51%*<br>(39% - 65%) | 51%*<br>(29% - 65%) | 47%*<br>(29% - 75%) | 37%*<br>(8% - 100%) | 62%*<br>(26% - 100%) | 28%*<br>(4% - 56%) | 60%*<br>(27% - 89%) |
| AOSIS | 7%*<br>(6% - 8%) | 4%*<br>(2% - 6%) | 3%*<br>(0% - 6%) | 0%<br>(0% - 51%) | 2%<br>(0% - 33%) | 0%<br>(0% - 6%) | 3%<br>(0% - 12%) |
| Indonesia | 7%*<br>(6% - 8%) | 4%*<br>(2% - 7%) | 3%*<br>(1% - 7%) | 0%<br>(0% - 51%) | 2%<br>(0% - 43%) | 0%<br>(0% - 6%) | 3%<br>(0% - 13%) |
| India | 11%*<br>(9% - 15%) | 10%*<br>(5% - 14%) | 7%*<br>(2% - 13%) | 3%<br>(0% - 100%) | 8%<br>(0% - 86%) | 4%<br>(0% - 16%) | 9%<br>(2% - 28%) |
| Japan | 14%*<br>(11% - 18%) | 13%*<br>(7% - 19%) | 9%*<br>(3% - 16%) | 5%<br>(0% - 100%) | 10%<br>(0% - 86%) | 7%<br>(0% - 21%) | 11%<br>(3% - 32%) |
| Russia | 20%*<br>(15% - 28%) | 21%*<br>(11% - 30%) | 16%*<br>(7% - 26%) | 10%<br>(0% - 100%) | 18%<br>(3% - 100%) | 12%<br>(0% - 30%) | 19%*<br>(7% - 46%) |
| EU27+UK | 47%*<br>(36% - 61%) | 48%*<br>(26% - 62%) | 42%*<br>(25% - 72%) | 33%<br>(6% - 100%) | 55%*<br>(22% - 100%) | 26%*<br>(4% - 51%) | 54%*<br>(24% - 86%) |
| Australia | 7%*<br>(6% - 9%) | 5%*<br>(3% - 8%) | 4%*<br>(1% - 8%) | 0%<br>(0% - 67%) | 3%<br>(0% - 43%) | 0%<br>(0% - 8%) | 4%<br>(0% - 16%) |

**Table S6.** Percentage of events with coral bleaching greater than 20% which are attributed to the emissions of countries and companies from 1985–2024. Columns show the fraction of bleaching events at coral reef sites globally and within specific countries or country blocs that can be attributed to emissions from specific companies and countries (listed in the rows). Values are medians and uncertainty estimates are 95% confidence intervals. Asterisks (*) denote emitters for which the attributed fraction is positive across the entire bootstrap distribution (1st percentile > 0%), indicating a virtually certain contribution to bleaching in that region. Top 10 refers to the top 10 major carbon emitting companies as defined in the Carbon Majors Database, AOSIS refers to the intergovernmental coalition the Alliance of Small Island States, and EU27+UK refers to the 27 countries of the European Union and the United Kingdom. *N* denotes the number of coral reef sites present in the data for each country or bloc.

| Scenario | Global (N=4471) | AOSIS (N=311) | Indonesia (N=89) | Japan (N=21) | China (N=65) | USA (N=25) | Australia (N=154) |
| --- | --- | --- | --- | --- | --- | --- | --- |
| All anthropogenic | 99%*<br>(94% - 100%) | 100%*<br>(94% - 100%) | 100%*<br>(97% - 100%) | 100%*<br>(90% - 100%) | 100%*<br>(100% - 100%) | 100%*<br>(92% - 100%) | 100%*<br>(99% - 100%) |
| All carbon majors | 91%*<br>(76% - 100%) | 86%*<br>(59% - 100%) | 100%*<br>(83% - 100%) | 100%*<br>(68% - 100%) | 100%*<br>(100% - 100%) | 100%*<br>(39% - 100%) | 100%*<br>(85% - 100%) |
| Saudi Aramco | 16%*<br>(12% - 25%) | 9%*<br>(3% - 19%) | 14%*<br>(3% - 37%) | 17%<br>(0% - 100%) | 27%<br>(0% - 100%) | 4%<br>(0% - 67%) | 24%<br>(0% - 60%) |
| ExxonMobil | 15%*<br>(11% - 23%) | 8%*<br>(3% - 18%) | 12%*<br>(3% - 35%) | 13%<br>(0% - 100%) | 18%<br>(0% - 100%) | 0%<br>(0% - 50%) | 20%<br>(0% - 56%) |
| Chevron | 16%*<br>(11% - 24%) | 9%*<br>(3% - 19%) | 13%*<br>(3% - 36%) | 13%<br>(0% - 100%) | 19%<br>(0% - 100%) | 0%<br>(0% - 59%) | 21%<br>(0% - 59%) |
| Holcim Group | 7%*<br>(4% - 10%) | 3%*<br>(0% - 5%) | 3%<br>(0% - 12%) | 0%<br>(0% - 25%) | 0%<br>(0% - 43%) | 0%<br>(0% - 6%) | 3%<br>(0% - 23%) |
| Top 10 | 60%*<br>(46% - 79%) | 45%*<br>(23% - 77%) | 76%*<br>(50% - 100%) | 83%*<br>(17% - 100%) | 100%*<br>(67% - 100%) | 32%*<br>(7% - 100%) | 87%<br>(0% - 100%) |
| Gazprom | 16%*<br>(11% - 24%) | 9%*<br>(3% - 19%) | 13%*<br>(3% - 36%) | 14%<br>(0% - 100%) | 24%<br>(0% - 100%) | 0%<br>(0% - 60%) | 22%<br>(0% - 60%) |
| National Iranian Oil Company | 13%*<br>(9% - 19%) | 7%*<br>(2% - 14%) | 9%*<br>(2% - 27%) | 9%<br>(0% - 94%) | 12%<br>(0% - 100%) | 0%<br>(0% - 50%) | 16%<br>(0% - 50%) |
| BP | 13%*<br>(9% - 20%) | 7%*<br>(2% - 15%) | 9%*<br>(2% - 29%) | 8%<br>(0% - 80%) | 12%<br>(0% - 100%) | 0%<br>(0% - 50%) | 16%<br>(0% - 50%) |
| China | 36%*<br>(26% - 50%) | 21%*<br>(11% - 51%) | 47%*<br>(15% - 76%) | 50%<br>(0% - 100%) | 87%*<br>(33% - 100%) | 11%<br>(0% - 100%) | 54%<br>(0% - 88%) |
| USA | 54%*<br>(40% - 72%) | 38%*<br>(20% - 71%) | 72%*<br>(41% - 100%) | 75%<br>(6% - 100%) | 99%*<br>(54% - 100%) | 21%*<br>(6% - 100%) | 81%<br>(0% - 100%) |
| AOSIS | 7%*<br>(4% - 11%) | 3%*<br>(0% - 6%) | 3%<br>(0% - 12%) | 0%<br>(0% - 29%) | 1%<br>(0% - 57%) | 0%<br>(0% - 8%) | 3%<br>(0% - 25%) |
| Indonesia | 8%*<br>(5% - 11%) | 3%*<br>(1% - 7%) | 3%<br>(0% - 13%) | 0%<br>(0% - 50%) | 3%<br>(0% - 86%) | 0%<br>(0% - 10%) | 5%<br>(0% - 25%) |
| India | 13%*<br>(10% - 20%) | 7%*<br>(2% - 15%) | 9%*<br>(2% - 27%) | 13%<br>(0% - 100%) | 15%<br>(0% - 100%) | 0%<br>(0% - 59%) | 17%<br>(0% - 50%) |
| Japan | 15%*<br>(11% - 23%) | 9%*<br>(3% - 19%) | 13%*<br>(3% - 36%) | 14%<br>(0% - 100%) | 21%<br>(0% - 100%) | 0%<br>(0% - 60%) | 21%<br>(0% - 58%) |
| Russia | 23%*<br>(16% - 33%) | 14%*<br>(6% - 29%) | 27%*<br>(6% - 56%) | 24%<br>(0% - 100%) | 41%<br>(0% - 100%) | 6%<br>(0% - 80%) | 35%<br>(0% - 75%) |
| EU27+UK | 50%*<br>(36% - 68%) | 34%*<br>(17% - 67%) | 69%*<br>(29% - 100%) | 64%<br>(6% - 100%) | 96%*<br>(47% - 100%) | 18%*<br>(5% - 100%) | 76%<br>(0% - 100%) |
| Australia | 8%*<br>(6% - 12%) | 4%*<br>(1% - 8%) | 3%<br>(0% - 15%) | 0%<br>(0% - 50%) | 4%<br>(0% - 90%) | 0%<br>(0% - 14%) | 6%<br>(0% - 30%) |

**Table S7.** Percentage of events with coral bleaching greater than 10% which are attributed to the emissions of countries and companies for 2024. Columns show the fraction of bleaching events at coral reef sites globally and within specific countries or country blocs that can be attributed to emissions from specific companies and countries (listed in the rows). Values are medians and uncertainty estimates are 95% confidence intervals. Asterisks (*) denote emitters for which the attributed fraction is positive across the entire bootstrap distribution (1st percentile > 0%), indicating a virtually certain contribution to bleaching in that region. Top 10 refers to the top 10 major carbon emitting companies as defined in the Carbon Majors Database, AOSIS refers to the intergovernmental coalition the Alliance of Small Island

| Scenario | Global (N=4471) | AOSIS (N=311) | Indonesia (N=89) | Japan (N=21) | China (N=65) | USA (N=25) | Australia (N=154) |
| --- | --- | --- | --- | --- | --- | --- | --- |
| **All anthropogenic** | **97%\***<br>(79% - 100%) | **99%\***<br>(79% - 100%) | **100%\***<br>(51% - 100%) | **100%\***<br>(56% - 100%) | **100%\***<br>(100% - 100%) | **100%\***<br>(100% - 100%) | **100%\***<br>(73% - 100%) |
| **All carbon majors** | 76%*<br>(51% - 95%) | 74%*<br>(54% - 100%) | 99%*<br>(53% - 100%) | 100%<br>(0% - 100%) | 100%<br>(0% - 100%) | 33%<br>(0% - 100%) | 100%*<br>(49% - 100%) |
| **Saudi Aramco** | 7%*<br>(3% - 15%) | 7%*<br>(2% - 14%) | 0%<br>(0% - 27%) | 0%<br>(0% - 100%) | 0%<br>(0% - 86%) | 0%<br>(0% - 15%) | 12%<br>(0% - 79%) |
| **ExxonMobil** | 6%*<br>(3% - 12%) | 6%*<br>(1% - 12%) | 0%<br>(0% - 22%) | 0%<br>(0% - 100%) | 0%<br>(0% - 86%) | 0%<br>(0% - 11%) | 9%<br>(0% - 74%) |
| **Chevron** | 6%*<br>(3% - 12%) | 6%*<br>(1% - 12%) | 0%<br>(0% - 23%) | 0%<br>(0% - 100%) | 0%<br>(0% - 86%) | 0%<br>(0% - 12%) | 10%<br>(0% - 74%) |
| **Holcim Group** | 3%*<br>(2% - 6%) | 2%*<br>(0% - 5%) | 0%<br>(0% - 11%) | 0%<br>(0% - 50%) | 0%<br>(0% - 3%) | 0%<br>(0% - 0%) | 2%<br>(0% - 19%) |
| **Top 10** | 39%*<br>(21% - 58%) | 40%*<br>(21% - 56%) | 51%*<br>(11% - 98%) | 15%<br>(0% - 100%) | 0%<br>(0% - 100%) | 10%<br>(0% - 36%) | 77%*<br>(27% - 100%) |
| **Gazprom** | 6%*<br>(3% - 14%) | 7%*<br>(2% - 14%) | 0%<br>(0% - 26%) | 0%<br>(0% - 100%) | 0%<br>(0% - 86%) | 0%<br>(0% - 14%) | 11%<br>(0% - 78%) |
| **National Iranian Oil Company** | 5%*<br>(3% - 11%) | 5%*<br>(1% - 11%) | 0%<br>(0% - 20%) | 0%<br>(0% - 84%) | 0%<br>(0% - 86%) | 0%<br>(0% - 10%) | 8%<br>(0% - 61%) |
| **BP** | 5%*<br>(3% - 10%) | 5%*<br>(1% - 10%) | 0%<br>(0% - 20%) | 0%<br>(0% - 67%) | 0%<br>(0% - 79%) | 0%<br>(0% - 10%) | 6%<br>(0% - 56%) |
| **China** | 21%*<br>(10% - 35%) | 21%*<br>(9% - 36%) | 17%<br>(0% - 73%) | 0%<br>(0% - 100%) | 0%<br>(0% - 100%) | 10%<br>(0% - 36%) | 44%*<br>(11% - 98%) |
| **USA** | 32%*<br>(15% - 49%) | 32%*<br>(16% - 49%) | 41%*<br>(6% - 92%) | 8%<br>(0% - 100%) | 0%<br>(0% - 100%) | 10%<br>(0% - 36%) | 65%*<br>(21% - 100%) |
| **AOSIS** | 3%*<br>(2% - 6%) | 2%*<br>(0% - 5%) | 0%<br>(0% - 12%) | 0%<br>(0% - 51%) | 0%<br>(0% - 12%) | 0%<br>(0% - 0%) | 2%<br>(0% - 26%) |
| **Indonesia** | 3%*<br>(2% - 7%) | 2%*<br>(0% - 6%) | 0%<br>(0% - 13%) | 0%<br>(0% - 51%) | 0%<br>(0% - 37%) | 0%<br>(0% - 0%) | 2%<br>(0% - 32%) |
| **India** | 6%*<br>(3% - 13%) | 6%*<br>(1% - 12%) | 0%<br>(0% - 23%) | 0%<br>(0% - 100%) | 0%<br>(0% - 86%) | 0%<br>(0% - 13%) | 10%<br>(0% - 74%) |
| **Japan** | 6%*<br>(3% - 13%) | 7%*<br>(2% - 13%) | 0%<br>(0% - 25%) | 0%<br>(0% - 100%) | 0%<br>(0% - 86%) | 0%<br>(0% - 14%) | 11%<br>(0% - 75%) |
| **Russia** | 9%*<br>(4% - 20%) | 10%*<br>(3% - 20%) | 1%<br>(0% - 35%) | 0%<br>(0% - 100%) | 0%<br>(0% - 100%) | 0%<br>(0% - 29%) | 16%<br>(0% - 85%) |
| **EU27+UK** | 28%*<br>(13% - 44%) | 28%*<br>(13% - 45%) | 37%<br>(3% - 90%) | 5%<br>(0% - 100%) | 0%<br>(0% - 100%) | 10%<br>(0% - 36%) | 61%*<br>(17% - 100%) |
| **Australia** | 3%*<br>(2% - 7%) | 3%*<br>(0% - 7%) | 0%<br>(0% - 13%) | 0%<br>(0% - 67%) | 0%<br>(0% - 37%) | 0%<br>(0% - 0%) | 2%<br>(0% - 35%) |

**Table S8.**
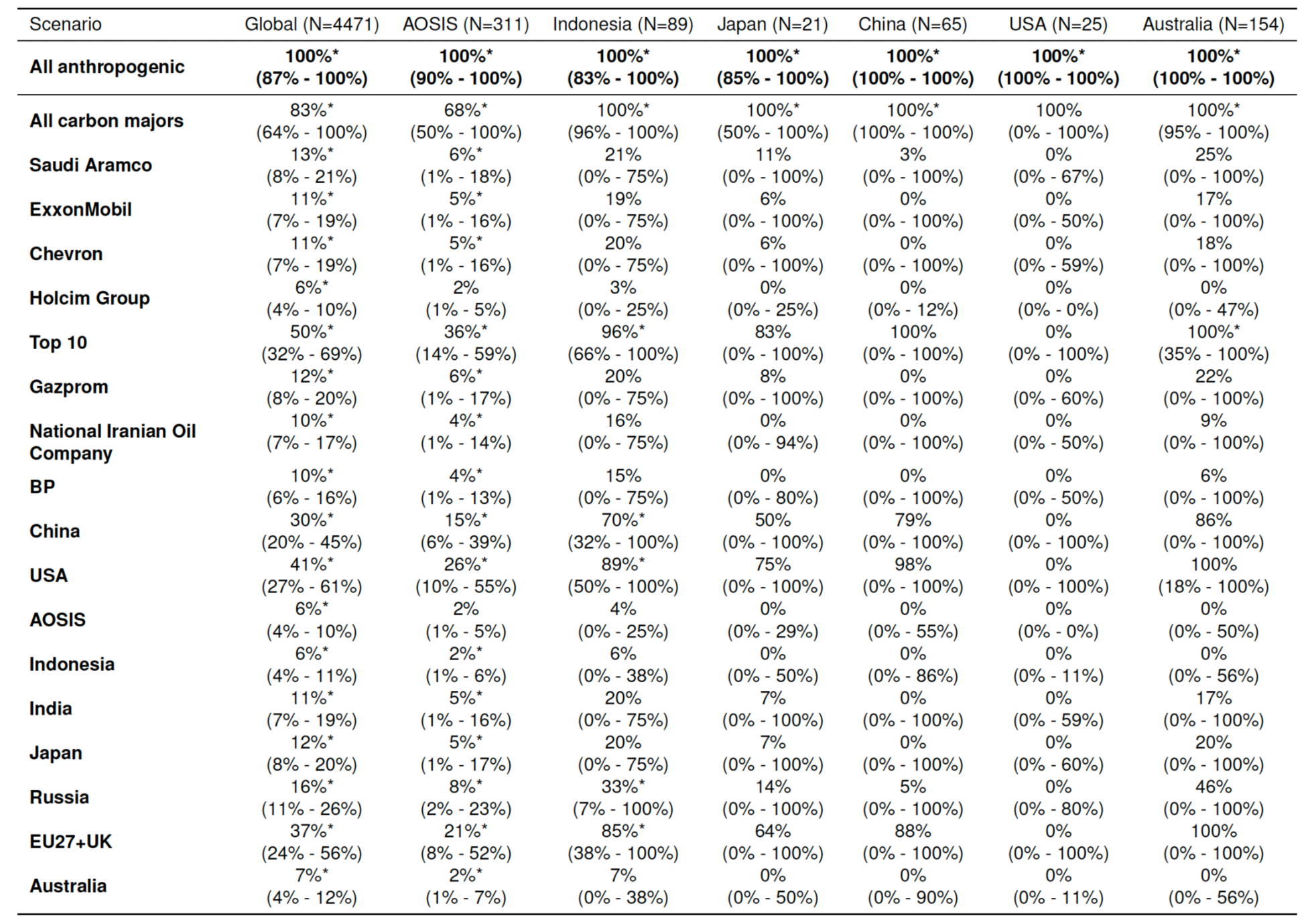
Percentage of events with coral bleaching greater than 20% which are attributed to the emissions of countries and companies for 2024. Columns show the fraction of bleaching events at coral reef sites globally and within specific countries or country blocs that can be attributed to emissions from specific companies and countries (listed in the rows). Values are medians and uncertainty estimates are 95% confidence intervals. Asterisks (*) denote emitters for which the attributed fraction is positive across the entire bootstrap distribution (1st percentile > 0%), indicating a virtually certain contribution to bleaching in that region. Top 10 refers to the top 10 major carbon emitting companies as defined in in the Carbon Majors Database, AOSIS refers to the intergovernmental coalition the Alliance of Small Island States, and EU27+UK refers to the 27 countries of the European Union and the United Kingdom. *N* denotes the number of coral reef sites present in the data for each country or bloc.

**Table S9.** Lag and lead effects of DHW on bleaching. All columns show results for panel regression models with percentage bleached as the dependent variable. All models include site, year and seasonality(ecoregion x month) fixed effects, with standard errors accounting for spatial autocorrelation using the Conley (200 km) method. Lag columns show DHW measured in the months following the bleaching survey; Lead columns show DHW measured in the months preceding it. Consistent with the definition of DHW as cumulative exposure over the past three months (see Methods), we see significant impacts of DHW exposure on bleaching at both lags and leads of the recorded bleaching event, with magnitudes and significance that decay to insignificant values at lags or leads greater than two months.

|  | *Dependent variable: Percent Bleached (%)* |  |  |  |  |  |  |  |  |
| --- | --- | --- | --- | --- | --- | --- | --- | --- | --- |
|  | Lag 3<br>(1) | Lag 2<br>(2) | Lag 1<br>(3) | No Lag or Lead<br>(4) | Lead 1<br>(5) | Lead 2<br>(6) | Lead 3<br>(7) | Lead 4<br>(8) | Lead 5<br>(9) |
| DHW Lag 3 | −0.45<br>(0.97) |  |  |  |  |  |  |  |  |
| DHW Lag 3 × \|Latitude\| | 0.00<br>(0.04) |  |  |  |  |  |  |  |  |
| DHW Lag 2 |  | 2.20*<br>(0.95) |  |  |  |  |  |  |  |
| DHW Lag 2 × \|Latitude\| |  | −0.08<br>(0.04) |  |  |  |  |  |  |  |
| DHW Lag 1 |  |  | 3.11**<br>(1.02) |  |  |  |  |  |  |
| DHW Lag 1 × \|Latitude\| |  |  | −0.11*<br>(0.04) |  |  |  |  |  |  |
| DHW |  |  |  | 3.82**<br>(1.09) |  |  |  |  |  |
| DHW × \|Latitude\| |  |  |  | −0.13**<br>(0.05) |  |  |  |  |  |
| DHW Lead 1 |  |  |  |  | 3.59**<br>(1.07) |  |  |  |  |
| DHW Lead 1 × \|Latitude\| |  |  |  |  | −0.12*<br>(0.05) |  |  |  |  |
| DHW Lead 2 |  |  |  |  |  | 0.49<br>(0.73) |  |  |  |
| DHW Lead 2 × \|Latitude\| |  |  |  |  |  | 0.01<br>(0.04) |  |  |  |
| DHW Lead 3 |  |  |  |  |  |  | −0.13<br>(0.56) |  |  |
| DHW Lead 3 × \|Latitude\| |  |  |  |  |  |  | 0.03<br>(0.04) |  |  |
| DHW Lead 4 |  |  |  |  |  |  |  | −0.01<br>(0.83) |  |
| DHW Lead 4 × \|Latitude\| |  |  |  |  |  |  |  | 0.02<br>(0.05) |  |
| DHW Lead 5 |  |  |  |  |  |  |  |  | 0.52<br>(1.32) |
| DHW Lead 5 × \|Latitude\| |  |  |  |  |  |  |  |  | 0.00<br>(0.08) |
| Observations | 6897 | 6897 | 6897 | 6897 | 6897 | 6897 | 6897 | 6897 | 6897 |
| $R^2$ | 0.551 | 0.559 | 0.570 | 0.582 | 0.579 | 0.553 | 0.551 | 0.550 | 0.552 |
| $R^2$ Within | 0.002 | 0.020 | 0.045 | 0.071 | 0.064 | 0.006 | 0.002 | 0.001 | 0.004 |
| BIC | 65593.6 | 65467.5 | 65291.0 | 65101.5 | 65150.9 | 65567.8 | 65597.1 | 65602.4 | 65578.9 |

\* $p < 0.05$ , \*\* $p < 0.01$

